# Distribution Shapes Govern the Discovery of Predictive Models for Gene Regulation

**DOI:** 10.1101/154401

**Authors:** Brian E. Munsky, Guoliang Li, Zachary R. Fox, Douglas P. Shepherd, Gregor Neuert

## Abstract

Despite substantial experimental and computational efforts, mechanistic modeling remains more predictive in engineering than in systems biology. The reason for this discrepancy is not fully understood. Although randomness and complexity of biological systems play roles in this concern, we hypothesize that significant and overlooked challenges arise due to specific features of single-molecule events that control crucial biological responses. Here we show that modern statistical tools to disentangle complexity and stochasticity, which assume normally distributed fluctuations or enormous datasets, don't apply to the discrete, positive, and non-symmetric distributions that characterize spatiotemporal mRNA fluctuations in single-cells. We demonstrate an alternate approach that fully captures discrete, non-normal effects within finite datasets. As an example, we integrate single-molecule measurements and these advanced computational analyses to explore Mitogen Activated Protein Kinase induction of multiple stress response genes. We discover and validate quantitatively precise, reproducible, and predictive understanding of diverse transcription regulation mechanisms, including gene activation, polymerase initiation, elongation, mRNA accumulation, spatial transport, and degradation. Our model-data integration approach extends to any discrete dynamic process with rare events and realistically limited data.

**Significance Statement:** Systems biology seeks to combine experiments with computation to predict complex biological behaviors. However, despite tremendous data and knowledge, most biological models make terrible predictions. By analyzing single-cell-single-molecule measurements of mRNA in yeast during stress response, we explore how prediction accuracy is controlled by experimental distributions shapes. We find that asymmetric data distributions, which arise in measurements of positive quantities, can cause standard modeling approaches to yield excellent fits but make meaningless predictions. We demonstrate advanced computational tools that solve this dilemma and achieve predictive understanding of many spatiotemporal mechanisms of transcription control including RNA polymerase initiation and elongation and mRNA accumulation, transport and decay. Our approach extends to any discrete dynamic process with rare events and realistically limited data.

---

The ultimate goal of modeling is to integrate quantitative data to understand, predict, or control complex processes. Useful models may be discovered through mechanistic or statistical approaches, but success is always limited by the quantity and quality of data and the rigor of comparison between models and experiments. These issues are largely solved in engineering, where computer analyses routinely enable the design of extraordinarily complex systems. Many would argue that predictive modeling in biology is far behind this capability due to limited experimental data, inescapable randomness or noise, and overwhelming biological complexity. These concerns have driven rapid single-cell experimental and computational advances, which have enabled measurement and modeling of individual biomolecules (i.e., DNA, RNA, and protein) in single cells with outstanding spatiotemporal resolution^1-12^. Such experiments have allowed the characterization of many intriguing aspects of biological complexity and variation^13^, while capturing these phenomena with stochastic gene regulation models has improved understanding of mechanisms and their parameters^14-18^.

Despite experimental and computational advances, most biological models still underperform expectations. While it is tempting to attribute this failure to “poor models” or “insufficient data,” an alternative and often overlooked explanation is that combinations of sufficient data and good models may fail because they haven't been integrated properly. Many standard engineering techniques exist to integrate models with continuous-valued data, but unlike most engineered systems, biological fluctuations are dominated by *discrete* events. A single molecule of DNA, RNA, or protein can change the fate of an organism^19-22^. The resulting positive and discrete distributions violate the most basic assumption of most model inference approaches (i.e., that measurement errors are continuous Gaussian random variables). Moreover, this violation is compounded by the fact that datasets for single-cell imaging and sequencing are usually too small to invoke the central limit theorem (CLT). Consequently, standard data-model integration procedures can fail dramatically. We hypothesize that more exact treatment of discrete biological fluctuations could solve the data-model integration dilemma and enable precise quantitative predictions.

To test this hypothesis, we explore the effects of experimental data distribution shapes on the uncertainty, bias, and resulting predictive capabilities of single-cell gene regulation models. We examine the evolutionary conserved Stress Activated Protein Kinase (p38 / Hog1 SAPK) signal transduction pathway (Figs. 1 and S1), and we quantify its control of transcription mechanisms including RNA polymerase transcription initiation and elongation on target genes as well as mature mRNA export and degradation in *Saccharomyces cerevisiae* during adaptation to hyper-osmotic shock (Fig. 1A)^23^. We quantify the number of individual mRNA primary transcripts at the site of transcription, in the nucleus, and in the cytoplasm for multiple genes using singlemolecule fluorescence *in situ* hybridization (smFISH) (Fig. 1B,C)^1,2^. We collect high-resolution data from more than 65,000 cells, and we quantify single-cell spatiotemporal mRNA distributions that are demonstrably non-normal and non-symmetric (Figs. S2-S3). For such distributions, huge data sets would be needed to justify application of the CLT as we discuss below. We use computational analyses to integrate these data with a discrete stochastic spatiotemporal model^24^ (Fig. 1A), and we show how different computational analyses of the *same experimental data* and *same models* can yield vastly different parameter biases and uncertainties which cause predictive accuracy to vary by many orders of magnitude (Fig. 1D).

**Figure 1.**
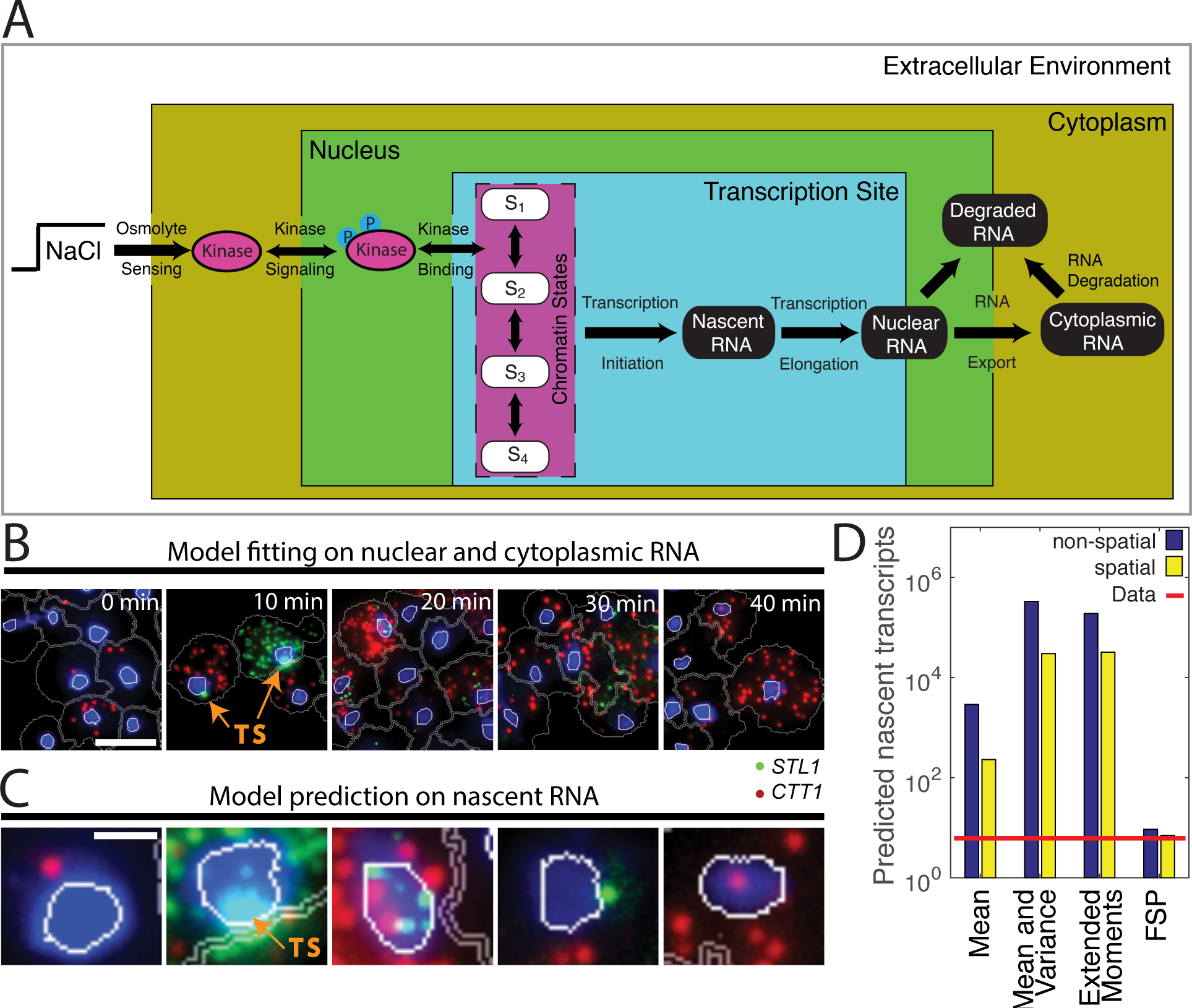
Discovering stochastic models to predict single-cell gene regulation. **(A)** Scope of the model to be identified, including quantitative analysis of MAPK induction and translocation, chromatin reorganization, polymerase initiation and elongation, and mRNA transcription, export, nuclear and cytoplasmic degradation. **(B)** Collection of single-cell spatiotemporal RNA transcription data to constrain the model (Fitting). Cytoplasmic and nuclear transcription quantification for expression of two mRNA species *(CTT1* in red and *STL1* in green). DAPI stained nucleus in blue. White line is the nuclear border, and the grey line is the cell boundary after automated segmentation. Representative images of cells exposed to 0.2M NaCl. 65454 cells in total have been imaged at 16 time points. Scale bar: 5μm. **(C)** Single-gene transcription site (TS) data used to validate the model (Prediction). Intensely bright spots within some cell nuclei are identified as transcription sites (TS). Scale bar: 1μm. **(D)** Validation of the model by comparing measured (solid red line) to predicted average number of nascent transcripts at active transcription sites using the same model and same data but under different modeling assumptions. Non-spatial analyses (blue) use the statistics (means, means and variances, or distributions) of the total number of RNA per cell. Spatial analyses (yellow) use the joint statistics of nuclear and cytoplasmic number of RNA per cell.

We discover that standard single-cell modeling approaches, which assume continuous and normally distributed fluctuations *or* enough data to invoke the CLT^24^, are not always valid to interpret finite datasets for single-cell transcription responses. We find that these standard approaches can yield surprising errors and poor predictions (Fig. 1D), especially when mRNA expression is very low. In contrast, we show that improved computational analyses of full single-cell RNA distributions can yield far more precisely constrained, less biased, and more reproducible models (Fig. 1D). We also discover new and valuable information contained in the intracellular spatial locations of RNA (Figs. 1B, S4-S5), enabling quantitative predictions for novel dynamics of gene regulation, including transcription initiation and elongation rates, fractions of actively transcribing cells, and the average number and distribution of polymerases per active transcription site, which could not otherwise be measured simultaneously in endogenous cell populations (Figs. 1D, 4).

**Figure 4.**
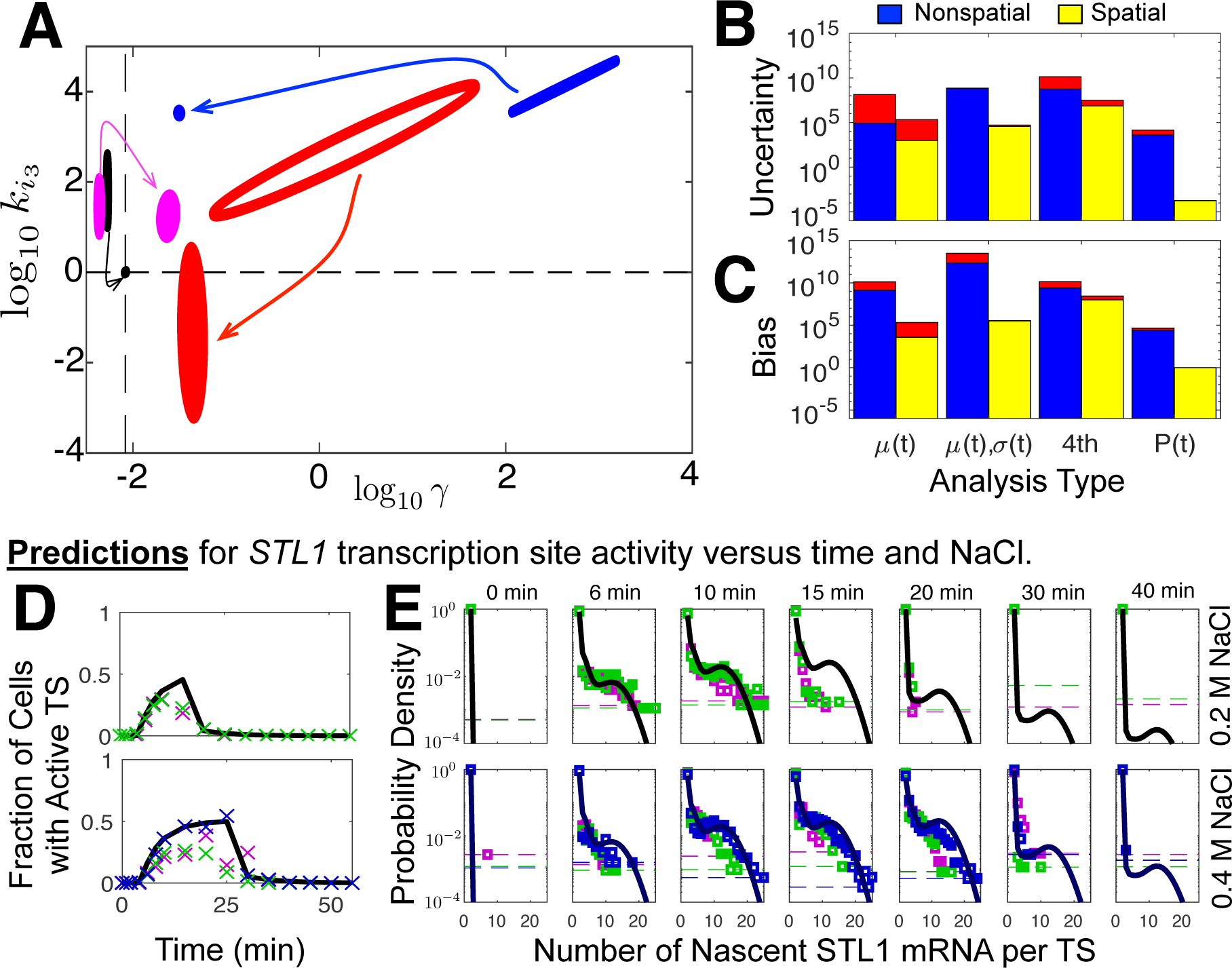
Stochastic and spatial fluctuation information improve parameter estimation to enable precise quantitative predictions. **(A)** Ninety percent confidence ellipses for the degradation rate (γ) and the maximal transcription initiation rate (*k_i3_*) using the means only (μ(t), red), means and variances (μ(t),Σ(t) blue), extended moment analyses (4th, magenta), or the full FSP distributions (**P**(t), black). Arrows show the effect of adding spatial information to the analyses. The dashed black lines show the fit parameters for the spatial FSP *STL1* model. **(B)** Total parameter uncertainty and **(C)** bias for the four analyses using non-spatial (blue) and spatial (yellow) analyses. The red regions show the difference between independent MHA chains. **(D)** Predicted (black) and measured (magenta, green and blue crosses) fractions of cells with active *STL1* TS versus time at 0.2M (top) NaCl and 0.4M (bottom) NaCl osmotic shock. **(E)** Predicted (black) and measured (magenta, green and blue) distributions of nascent *STL1* mRNA per TS at different times following 0.2M (top) NaCl and 0.4M (bottom) NaCl osmotic shock. Magenta, green and blue horizontal lines correspond to the minimum detection limit (1/N_c_, where N_c_ is the number of cells measured at that time for the corresponding biological replica). All predictions are made using parameters estimated previously from the mature, spatial mRNA distributions.

## Results

Under osmotic stress, the high osmolarity glycerol kinase, Hog1, is phosphorylated and translocated to the nucleus, where it activates several hundred genes^23^. For two of these genes (*STL1*, a glycerol proton symporter of the plasma membrane and *CTT1,* the Cytosolic catalase T), we quantified transcription at single-molecule and single-cell resolution (Figs. 1B,C, S2, and S3), at temporal resolutions of one to five minutes, at two osmotic stress conditions (0.2M and 0.4M NaCl), and in multiple biological replicas. We built histograms to quantify the marginal and joint distributions of the nuclear and cytoplasmic mRNA (Figs. 2D,E and S2-S5).

**Figure 2.**
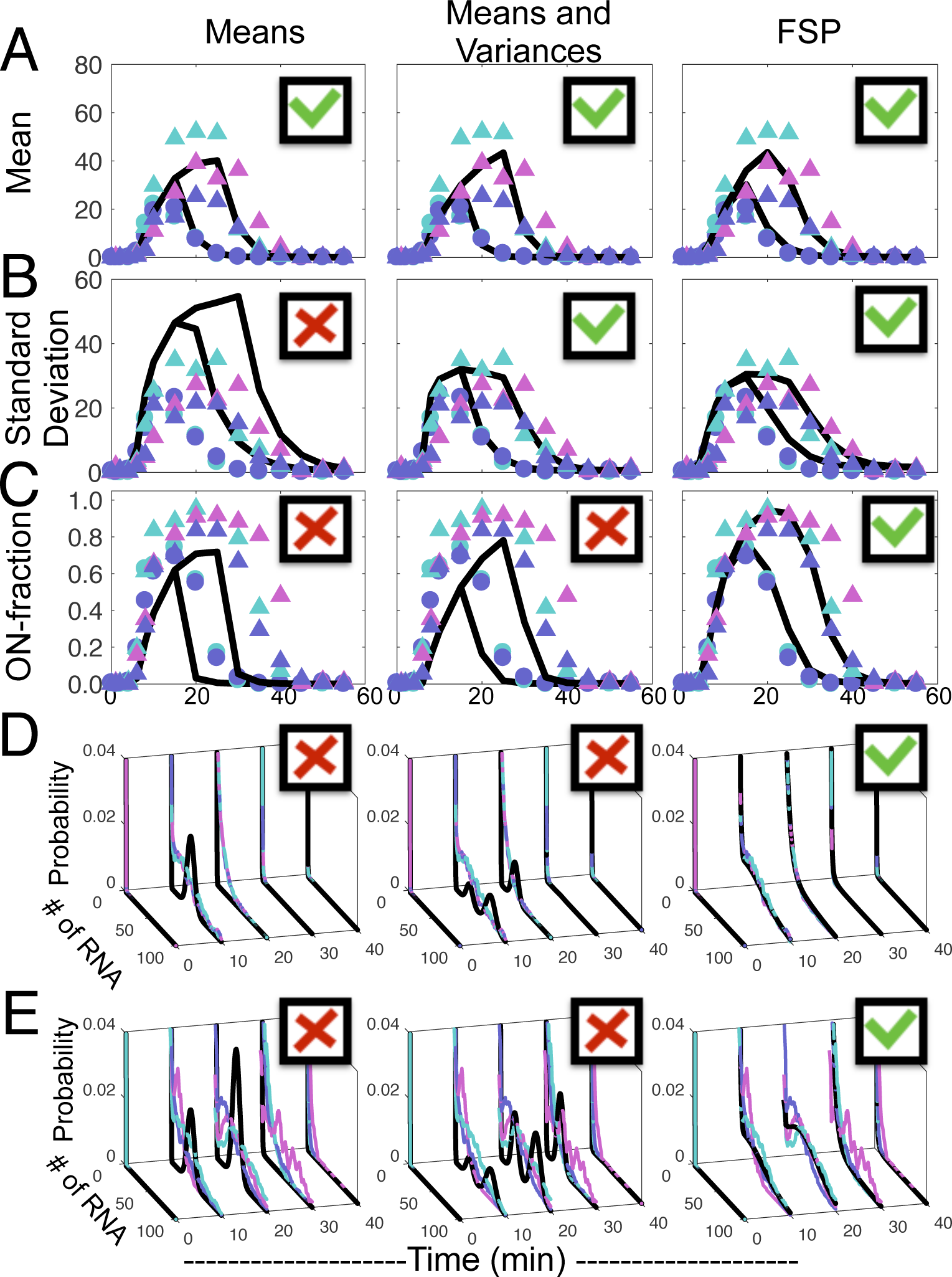
Different computational analyses result in matches to different data statistics. **(A)** Mean number, **(B)** standard deviation, **(C)** ON-fraction (cells with >3 mRNA), and **(D,E)** distributions of *STL1* mRNA copy number. In panels **A-C**, data for 0.2M NaCl (circles, two biological replica) and 0.4M NaCl (triangles, three biological replica) are shown in magenta, cyan, and violet, and model results are shown in black. Temporal distributions are measured at **(D)** 0.2M NaCl and **(E)** 0.4M NaCl (cyan, violet, and magenta are biological replica). The left column corresponds to the best fit to the measured mean; the center column corresponds to the best fit to the measured means and variances; and the right column corresponds to the best fit to the full measured distributions.

We extended a bursting gene expression model^14,25^ to account for transcriptional regulation and spatial localization of mRNA (Fig. 1A)^24^. We considered four approaches to fit this model to gene transcription data: First, we used exact analyses of the first moments (i.e., population means) of mRNA levels as functions of time. Second, we added exact analyses of the second moments (i.e., variances and covariances). Third, we extended the moments analyses to include the third and fourth moments. Finally, we used the finite state projection (FSP^26^) approach to compute the full joint probability distributions for nuclear and cytoplasmic mRNA. *All four approaches provide exact solutions of the same model*, but at different levels of statistical detail^24^. We used each analysis to compute the likelihood that the measured mRNA data would match the model, and we maximized this likelihood^24^. As was the case for previous studies^17,27^, we note that the moments-based likelihood computations assume either normally distributed deviations (first and second methods) *or* sufficiently large sample sizes such that the moments can be captured by a normal distribution as guaranteed by the CLT (third method)^24^. In contrast, the FSP approach (fourth method) makes no assumptions on the distribution shape and has no requirement for large sample sizes.

### Different exact analyses of the same model and same data yield dramatically different results

The four likelihood definitions were maximized by different parameter combinations (Tables S3 and S4), and the fit and prediction results are compared to the measured mean, variance, ON-fraction (i.e., fraction of cells with more than 3 mRNA / cell), and distributions versus time for *STL1* and *CTT1* (Figs. 2, S2, and S3). The different analyses used the same model, and they were fit to the exact same experimental data, but they yielded dramatically different results. When identified using the average mRNA dynamics (Fig. 2A, left), the model failed to match the variance, ON-fractions, or distributions of the process (Fig. 2B-E, left). Fitting the response means and variances simultaneously (Fig. 2A-B, center) failed to predict the ON-fractions or probability distributions (Fig. 2C-E, center). In contrast, parameter estimation using the full probability distribution (Fig. 2, right column and Figs. S2 and S3) matched all measured statistics. Importantly, key parameters identified using the FSP approach agree well with previous studies^18^, such as *STL1* and *CTT1* mRNA transcription and degradation rates. This agreement indicates strong reproducibility of both experiments and analyses (Tables S3 and S4) and provides more confident predictions for new transcriptional mechanisms as discussed below. In contrast, the moment-based analyses overestimated these rates by multiple orders of magnitude.

### Standard modeling identification procedures fail due to bias in moment estimation

We considered three explanations for the failure of moment-based parameter estimation approaches: (i) the model parameters could be unidentifiable from the considered moments; (ii) the parameters could be too weakly constrained by those moments; or (iii) the moments analyses could have introduced systematic biases due to a failure of the CLT. To eliminate the first explanation, we computed the Fisher Information Matrix (FIM) defined by the moments-based analyses^24^. Because the computed FIM has full rank, we conclude that the model should be identifiable. If the second explanation were true (i.e., if the moments-analyses had produced weakly constrained models), then changing the parameters to those selected by the FSP analysis should have only a small effect on the moment-based likelihood. In such a case, the FSP parameters would lie within large parameter confidence intervals identified by the moments-based analyses. However, using the experimental *STL1* data, we computed that the FSP parameter set was 10^2,750^ less likely to have been discovered using means, 10^14,500^ less likely to have been discovered using means and variances, and 10^664^ less likely to have been discovered using the extended moments analysis (Table S5). Thus, we conclude that failure of the moments-based analyses to match the distributions in Figs. 2, S2, and S3 cannot be explained by uncertainty alone.

To test the third explanation for parameter estimation failure (i.e., systematic bias), we used the FSP parameters and generated simulated data for the mean (Fig. 3A), standard deviation (Fig. 3B), and the ON-fraction (Fig. 3C) versus time for *STL1* mRNA under an osmotic shock of 0.2M NaCl. As shown in Fig. 3A,B, the *median* of the simulated data sets (magenta) matches the experimental data (red and cyan) at all times, but at later times (>20 minutes) both are consistently less than the theoretical values (black). This mismatch is due to finite sampling from asymmetric distributions especially at later time points (Fig. 3D). The Gaussian assumption applied to the first two moments analyses^24^, which does not account for asymmetry, imposes narrow and nearly symmetric likelihood functions for the sample mean and sample variance (cyan lines in Fig. 3E,F, respectively). These moment-based likelihood functions are inconsistent with the actual sample statistic distributions (Fig. 3F, compare cyan and black lines). Because the mRNA distributions are very broad at late time points (Fig. 3D), one would need to measure 10^5^ or 10^7^ cells to estimate the variance within 10% or 1%, respectively (Fig. 3G). Furthermore, because the mRNA distributions are highly asymmetric, measurements are likely to repeatedly underestimate the summary statistics. As a result, the moment-based likelihood functions were deleteriously overfit to underestimated mRNA expression at late time points, and resulted in excessively confident overestimation of the mRNA degradation rate (Table S3). In principle, the extended moments analysis could better capture the correct likelihood function for the sample statistics (Figs. 3E,F magenta lines), but this requires *a prior* knowledge of the exact third and fourth moments. In practice, because those higher order statistics must be estimated using the model, the resulting extended moments analysis was less biased but more uncertain.

**Figure 3.**
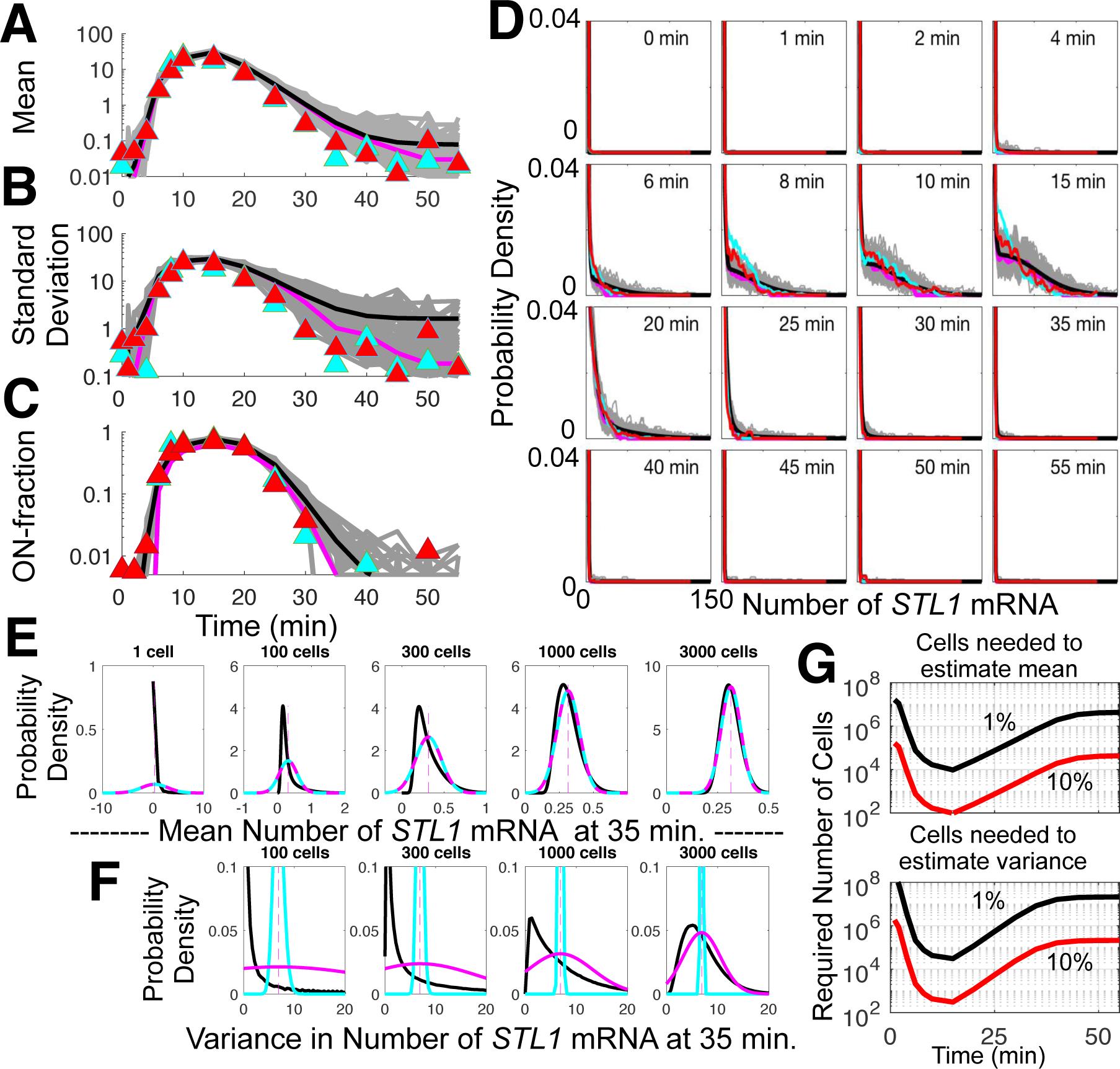
Discrete positive distributions introduce uncertainty and bias in parameter estimation. **(A)** Mean, **(B)** standard deviation, **(C)** ON-fraction, and **(D)** full distributions of *STL1* mRNA versus time for an osmotic shock of 0.2M NaCl applied at time *t* = 0. Theoretical values are in black, representative simulated samples of 200 cells each are in gray, median statistics of the simulated samples are in magenta; and experimental biological replica data are in red and cyan. **(E,F)** Expected distribution of sample mean **(E)** and sample variance **(F)** for *STL1* at 35 minutes computed using the CLT using Gaussian approximation (cyan), an extended moment analysis (magenta), or exact sampling from the FSP (black) for population sizes of 1, 100, 300, 1000, and 3000 cells. **(G)** Expected number of cells required to estimate the mean (top) or variance (bottom) within standard errors of 10% (red) or 1% (black). The dependence on time is due to the changing distribution shapes shown in Panel **D**.

### Full distribution analyses substantially reduce model uncertainty and bias

To confirm the tradeoff between uncertainty and bias, we applied the Metropolis Hastings algorithm (MHA) to analyze parameter variation for the different likelihood functions and to estimate parameter uncertainty and bias (Fig. 4A-C)^24^. Comparing the parameter variations for the transcription initiation rate, *k*_*i*3_, and the mRNA degradation rate, *γ*, illustrates that extending the analysis from the means to means and variances can affect the parameter identification bias much more than the parameter uncertainty (Fig. 4A). Moreover, this effect is not always advantageous; inclusion of variances in the analysis led to substantially increased parameter bias for *STL1* (compare red and blues ellipses in Fig. 4A,C and see Fig. S9) and relatively little change for *CTT1* (Figs. S10 and S11). In contrast, analyses using the FSP consistently reduced both uncertainty and bias for both *STL1* and *CTT1* analyses (Figs. 4A-C and S9-S11).

### Using spatial fluctuations improves model identification

Having established that different stochastic fluctuation analyses attain different levels of uncertainty and bias, we asked if more information could be extracted from spatially-resolved data. Using a nuclear stain, we quantified the numbers of *STL1* and *CTT1* mRNA in the nucleus and cytoplasm^24^ (Fig. 1B). We then extended the model and our analyses to consider the joint cytoplasmic and nuclear mRNA distributions (Fig. S4 and S5). From these analyses, we observed that spatial data reduced parameter bias for the models, despite the addition of new parameters and model complexity (Fig. 4A-C and S9-11).

### Measuring and predicting transcription site dynamics

We next explored how well the identified models could be used to predict the elongation dynamics of nascent mRNA at individual *STL1* or *CTT1* transcription sites (TS, Fig. 1B,C). We quantified the TS intensity for *CTT1*, and we used an extended FSP model for *CTT1* regulation to estimate the Polymerase II elongation rate to be 63±13 nt/s^24^, a value consistent with published rates of 14-61 nt/s^28,29^. We assumed an identical rate for the *STL1* gene, and we used the FSP model for *STL1* gene regulation to predict the *STL1* TS activity (Figs. 1D, 4D). The spatial (non-spatial) FSP model predicts an average of 7.0 (9.3) full length *STL1* mRNA per active TS, a value that matches well to our measured value of 4.2-7.5 *STL1* mRNA per active TS. However, predictions using parameters identified from moments-based analyses were incorrect by several orders of magnitude (Fig. 1D). In addition to predicting the average number of nascent mRNA per active TS, the FSP model also accurately predicts the fraction of cells that have an active *STL1* TS versus time as well as the distribution of nascent mRNA per TS (Figs. 4E).

## Discussion

Integrating stochastic models and single-molecule and single-cell experiments can provide a wealth of information about gene regulatory dynamics^14^. In previous work, we discussed the importance to *choose the right model* to match the single-cell fluctuation information and achieve predictive understanding^18^. Here we have shown how important it is to *choose the right computational analysis* with which to analyze single-cell data. We showed how model identification based solely upon average behaviors can lead to substantial parameter uncertainty and bias, potentially resulting in poor predictive power (Figs. 1-4). We showed how single-molecule experiments often yield discrete, asymmetric distributions that are demonstrably non-Gaussian (Figs. 2D,E, 3, S2, and S3), and how model extensions to include hard-to-measure variances and covariances may exacerbate biases (Fig. 4C) leading to greatly diminished predictive power (Figs. 1D). By taking into account the full distribution shapes, one can correct these deleterious effects and obtain parameter estimates and predictions that are improved by many orders of magnitude, even when applied to the same model and same data (Figs. 1D, 2, 4). We stress that this concern occurs even for models for which exact equations are known and solvable for the statistical moment dynamics. For more complex and nonlinear models, where approximate moment analyses are required, these effects are likely to be exacerbated further. This issue is expected to be even more relevant in mammalian systems, which exhibit greater bursting^1,2,21^ and for which data collection may be limited to smaller sizes (e.g., by increased image processing difficulties for complex cell shapes or by small numbers of cells, as available from an organ, a tissue from a biopsy, or for a rare cell type population).

Because most single-cell modeling investigations to date have used only means or means and variances from finite data sets to constrain models, it is not surprising that many biological models fail to realize predictive capabilities. Conversely, our full consideration of the single-molecule distributions enabled discovery of a comprehensive model that quantitatively captures transcription regulation with biologically realistic rates and interpretation for transcription initiation, transcription elongation, and mRNA export and nuclear and cytoplasmic mRNA degradation (Fig. 1A). We argue that the solution is not to collect increasingly massive amounts of data, but instead to develop computational tools that utilize the full, unbiased spatiotemporal distributions of single-cell fluctuations. By addressing the limitations of current approaches and relaxing requirements for normal distributions or large sample sizes, our approach should have general implications to improve mechanistic model identification for any discipline that is confronted with non-symmetric datasets and finite sample sizes.

## Methods and Materials

### Yeast strain and growth condition

*Saccharomyces cerevisiae BY4741* (*MATa; his3*Δ*1; leu2*Δ*0; met15*Δ*0; ura3*Δ*0*) was used for time-lapse microscopy and FISH experiments. Cells are grown in complete synthetic media (CSM, Formedia, UK) to an optical density (OD) of 0.5. Nuclear enrichment of YFP C-terminus tagged Hog1 was measured in single cells over time in response to osmotic stress^30,31^.

### Microscopy setup for time-lapse and single molecule RNA FISH imaging

Cells were imaged with a Nikon Ti Eclipse epifluorescent microscope equipped with perfect focus (Nikon), a 100x VC DIC lens (Nikon), fluorescent filters for YFP, DAPI, TMR and CY5 (Semrock), an X-cite 120 fluorescent light source (Excelitas), and an Orca Flash 4v2 CMOS camera (Hamamatsu). The microscope was controlled by Micro-Manager program^32^.

### Image acquisition for time-lapse and single molecule RNA FISH imaging

Constant focus mode live cell time-lapse microscopy was performed by taking bright field images every 10 s and YFP fluorescent images every minute. Single molecule RNA-FISH microscopy was performed as z-stacks of 200nm between images from fixed yeast cells for bright field, DAPI, TMR and CY5.

### Sample preparation for live cell time-lapse microscopy

Hog1-YFP tagged yeast cells are loaded into a flow chamber^33^. CSM or CSM with 0.2M or 0.4M NaCl was passed through the flow chamber using a syringe pump (New Era Pump Systems) at a pump rate of 0.1 ml/minute.

### Image analysis for time-lapse microscopy

The bright field and YFP images are background corrected and smoothed. The YFP images are then thresholded and converted to a binary image resulting in a nuclear marker for cell segmentation. The processed bright field and the binary YFP images were combined using morphological reconstruction, and cells were segmented with a watershed algorithm. After segmentation, the centroid of each cell was computed and tracked over time to generate single cell time trajectories. For each cell the Hog1 nuclear enrichment was then calculated as Hog1(t) = [(I_t_(t) - I_b_) / (I_w_(t) - I_b_)], with I_w_ (average per pixel fluorescent intensity of the whole cell), I_b_ (average per pixel fluorescent intensity of the camera background), and I_t_ (average per pixel fluorescent intensity of the 100 brightest fluorescent pixels). The single cell traces were smoothed and subtracted by the Hog1(t) signal on the beginning of the experiment. For cell volume measurements, volume change relative to the volume at the beginning of the experiment was calculated. The median and the average median distance (standard deviation of the median) from single cell time traces of Hog1-YFP and cell volume was computed. The final time-lapse microscopy data set consist of 246 (0.2M NaCl) and 167 (0.4M NaCl) cells. Each time course was measured in duplicates or triplicates.

### Sample preparation for single-molecule RNA-FISH

Yeast cells (OD = 0.5) was concentrated tenfold through a filter system, then exposed to NaCl concentration of 0.2 M and 0.4 M NaCl, and then fixed in 4 % formaldehyde at time points 0, 1, 2, 4, 6, 8, 10, 15, 20, 25, 30, 35, 40, 45, 50, 55 minutes after osmotic stress. After fixation, cells are spheroplasted with 2.5 mg/ml Zymolyase (US Biological) until cells turn from an opaque to a black color. Cells are stored in 70 % ethanol at +4C for at least 12 hours. Hybridization and RNA-FISH probe preparation conditions were applied as published^18^. Table S1 contains the RNA-FISH probes for *STL1* and *CTT1*. Each time course was measured in duplicate or triplicate.

### Image analysis for single-molecule RNA-FISH

Each DAPI image stack was maximum intensity projected, thresholded, converted into a binary image, and individual connected regions of biologically relevant size was segmented resulting in individually labeled nuclei. For each cell, the entire DAPI image stack was converted into a binary image stack using cell specific DAPI thresholds resulting in 3D thresholded nucleus and cytoplasm. The last 5 images of the bright field image stack were maximum projected to generate the cell outlines. This image was background corrected and combined with the threshold DAPI image using morphological reconstruction and cells were segmented with a watershed algorithm. To identify RNA spots, a fluorescent intensity threshold was determined for each dye, and each field of view imaged. For a specific time course experiment, the average fluorescent threshold was determined from the average of all the individual thresholds. After the thresholds had been determined each image in the stack was Gaussian filtered, filtered with a Laplacian of a Gaussian filter to detect punctuate in the image, converted into binary images using the previously determined threshold, and the regional maxima was determined within the image stack. A mask of segmented nucleus and cytoplasm was applied to the filtered RNA-FISH image stacks and the numbers of RNA spots were counted in each compartment for each cell in three dimensions. To determine the number of transcripts of each transcription site, we determined the total intensity of each RNA spot in the cell. We also fitted each RNA spot with a 2D Gaussian function to determine the RNA spot intensity and found very good agreement between the fitted and the total RNA spot intensities. The total RNA-FISH data set consists of a total of 65454 single cells (25511 at 0.2M NaCl and 39943 at 0.4M NaCl).

### RNA-FISH data analysis

For each time point, a distribution of mRNA molecules in single cells was determined as the marginal distribution of total *STL1* and *CTT1* mRNA or as the joint probability distribution of nuclear and cytoplasmic RNA. The distributions were also summarized in a binary data set of ON-cells and OFF-cells. Cells with three or more mRNA molecules were considered ON-cells for quantification of the ON-fraction. The population average was computed as the mean marginal cytoplasmic or nuclear RNA for each mRNA species. To quantify transcription site intensities, we recorded the intensity of all spots in the nuclei for all cells. The brightest nuclear spots of each cell were labeled as potential transcription sites, and were temporarily removed from the data set. We then computed the median intensity for the remaining nuclear spots, and we used this median value to define the equivalent mature nuclear mRNA fluorescence intensity. Next, we re-examined the brightest RNA spot in each nucleus (i.e., the potential transcription sites), and we computed the fluorescence intensity in units of mature mRNA intensities. Potential transcription sites that had intensities greater than the equivalent of two mature mRNA molecules were labeled as active transcription sites. This approach was applied to all cells from a single experiment. From this analysis, we computed the fraction of cells with an active transcription site as a function of time, the single-cell distributions of full length RNA transcripts per transcription site as a function of time, and the average number of full length RNA transcripts per transcription site as a function of time.

### Hog1-Kinase Model

To model the temporally-varying Hog1p localization signal, we adopt the model from^18^, and we fit parameters of this model to the measured Hog1p nuclear enrichment levels as functions of time, and osmolyte concentrations (see Table S2 and Fig. S1). This time-varying signal has been used as an input to the gene regulation models.

### Gene Regulation Model

To capture the spatial stochastic expression of *STL1* or *CTT1* mRNA, the four-state *Hog1p*-activated gene expression model identified in Ref. 18 was extended to account for spatial localization of mRNA in the nucleus or cytoplasm (see Fig. 1A). In total, there are 13 non-spatial or 15 spatial parameters in the model.

### Computation of Moments

Moment dynamics were analyzed using sets of coupled linear time-varying ordinary differential equations. Theses ODEs provide exact expressions for the dynamics of the model’s means, variances, covariances and higher moments^34^. Because the reaction rates are all linear, these moment equations are closed, and the moments can be computed with no assumption on the distribution shape.

### Computation of Full Distributions

Distributions were computed using the Finite State Projection approach^26^ to approximate the exact solution of the Chemical Master Equation.

### Computation of Moment-Based Likelihood Functions

To compare computed moments to measured experimental data required computation of the likelihood that the measured moments (e.g., means or covariance matrix) would have been observed by chance given that the model were correct. Three different approaches were derived to compute the likelihood of the observed moments given the model. First, to estimate the likelihood that the average data could come from the model, fluctuations were assumed to be Gaussian, with the model-generated means and the measured sample (co)variances. Second, to compute the likelihood to observe both the measured sample variance (non-spatial) or covariance matrix (spatial), the chi-squared distribution (non-spatial) or Wishart distribution (spatial) were used to approximate the likelihood to obtain the measured sample means and variances, given the model^35^. Third, the approach from Ref. 17 (based upon application of the central limit theorem) was used to approximate the joint likelihood to simultaneously measure the join sample means and sample covariance matrix in which the co-variance of the first two moments was assumed to depend upon the first four moments of the model at each condition and point in time.

### Computation of the Full Distribution Likelihood

The likelihood of the full distribution data was computed using the FSP approach, similar to the approach taken in Refs. 18,36.

### Parameter Searches to Maximize Likelihood

Iterative combinations of simplex based searches and genetic algorithm based searches were used to maximize likelihood functions. All parameters were defined to have positive values, and searches were conducted in logarithmic space. The searches are run multiple times from different starting parameter guesses leading to many tens of millions of function evaluations (>4 × 10^7^ evaluations of the means and moments analyses, >5 × 10^6^ evaluations of the non-spatial FSP distributions, and >5 × 10^5^ evaluations for the more computationally expensive analyses of extended moments and the spatial FSP distributions). Parallel fits were conducted on clusters of more than 128 processors at a time allowing for the consideration of several millions of model/parameter/experiment combinations per day.

### Quantification of Parameter Uncertainties

The Metropolis Hastings algorithm^37^ was used to quantify the parameter uncertainties. All MCMC parameter explorations were conducted in logarithmic parameter space, and all analyses (i.e., means, means and variances, or distributions, both spatial and non-spatial) used the same proposal distribution for the MCMC chains. This proposal distribution was a symmetric normal distribution (in logarithmic space) with a variance of 0.005 (also in logarithmic space). For each parameter proposal, a random selection of parameters was selected to change, where each parameter had a 50% chance to be perturbed. The first half of each chain was discarded as an MCMC burn-in period, and all chains were thinned by 90%. MCMC chains for the means and simpler moments analyses were run for MH chains with lengths of 1,000,000 parameter evaluations. MCMC analyses for the more computationally expensive extended moment analyses were run for at least 120,000 parameter evaluations. MCMC analyses for the non-spatial distribution analyses were run for at least 250,000 parameter evaluations. MCMC analyses for the most computationally expensive spatial distribution analyses were run for at least 15,000 parameter evaluations. To evaluate convergence of the MCMC analyses, multiple MCMC chains were run for each analysis. To illustrate the achieved convergence, Fig. S8 shows the similarity of distributions of likelihood values for two independent MCMC runs for each analysis and for *STL1* and *CTT1.* The total bias and total uncertainty, shown in Figs. 4B, 4C, S11B and S11C, were computed as described in the supplemental material.

### Predictions of Transcription Site Activity

Two different analyses were developed to predict transcription site (TS) activity: a simplified theoretical analysis of average active TS activity and an extended FSP analysis of distributions of polymerases on a given TS. In the *simplified analysis*, it was assumed than an *active* TS would correspond to one gene at steady state with the maximum transcription rate, *k*_i-max_. Under this assumption, the average number of elongating polymerases is given by <n_pol_> = *k*_i-max_τ_elong_= *k*_i-max_*L/k*_elong_. The average nascent mRNA was assumed to be half the length of a mature mRNA, and since smFISH probes were equally distributed along the length of the gene, the average nascent mRNA was assumed to exhibit half the brightness of a mature mRNA. An *extended FSP approach* similar to that in (38) was also used to compute the distribution for the number of polymerases at the TS. Under the assumption that each partially transcribed mRNA was at a random location in the gene, its effective intensity was approximated to be uniformly distributed between zero and the equivalent of one mature mRNA. The distribution of TS spot intensities with *N*_poly_ polymerases was found through the convolution of *N*_poly_ independent random variables, each with a uniform distribution between zero and one. The FSP analysis was confirmed using an adapted form of the Stochastic Simulation Algorithm^39^ but with a modification^40^ to allow for time varying reaction rates (Fig. S12). For both experimental and computational analyses, TS sites were labeled as ON if their predicted or measured intensities were greater than twice the intensity of a single mature mRNA.

### Identification of mRNA Elongation Rate

The transcription elongation rate was found by computing the TS intensity distribution for *CTT1* at each point in time for 0.2M and 0.4M NaCl osmotic shock using the previously identified parameters (Table S4) and one free constant to describe the average elongation rate, *k*_elong_. The probability that the observed distributions of *CTT1* TS intensities could have originated from this model was computed for all time points and conditions, and as a function of *k*_elong_. This likelihood was maximized for the different biological replicas and NaCl concentrations to determine the uncertainty in this parameter. The simplified theoretical model, which does not account for transitions between active and inactive periods, provided an *upper bound* on the *CTT1* elongation rates to be 91±9Nt/s. The more detailed spatial FSP approach determined the *CTT1* elongation rates to be 63±14Nt/s. For both cases, the uncertainty is given as the standard error of the mean using the five experimental replicas (two for 0.2M NaCl and three for 0.4M NaCl). The elongation rate was then fixed to be 63Nt/s, and this rate was used in conjunction with the previously identified parameters to predict the TS intensity distributions for *CTT1* and *STL1* as functions of time in both osmotic shock conditions (Figs. 1D, 4D,E and S11D-H).

### Codes

All computational results presented in this paper can reproduced using the provided Matlab codes. The codes also include a simple GUI interface to generate additional figures: to make predictions with changes to the identified parameters, to produce confidence ellipses for new combinations of parameters, etc.

## Acknowledgments

BM and ZF were funded by the W.M. Keck Foundation, DTRA FRCALL 12-3-2-0002 and CSU Startup Funds. GL and GN were funded by NIH DP2 GM11484901, NIH R01GM115892 and Vanderbilt Startup Funds. The authors like to thank Luis Aguilera, Anthony Weil, Alexander Thiemicke, Dustin Rogers, Benjamin Kesler and Rohit Venkat for comments on the manuscript.

## Contributions

BM and ZF designed and implemented the computational analyses. GL and GN designed and implemented the experimental analyses. BM, ZF, DS and GN wrote the manuscript.

## Supplementary Materials for

### 1 Materials and Methods – Experimental

#### 1.1 Yeast strain and growth conditon

*Saccharomyces cerevisiae BY4741* (*MATa; His3Δ1; leu2Δ0; met15Δ0; ura3Δ0*) was used for time-lapse microscopy and FISH experiments. Three days before the experiment, yeast cells from a stock of cells stored at -80°C were streaked out on a complete synthetic media plate (CSM, Formedia, UK). The day before the experiment, a single colony from the CSM plate was inoculated in 5 ml CSM medium (preculture). After 6-12h, the optical density (OD) of the pre-culture was measured and the cells were diluted into new CSM medium to reach an OD of 0.5 the next morning.To assay the nuclear enrichment of Hog1 in single cells over time in response to osmotic stress, a yellow-fluorescent protein (YFP) was tagged to the C-terminus of endogenous Hog1 in *BY4741* cells through homologous DNA recombination [1] [2].

#### 1.2 Microscopy setup for time-lapse and single molecule RNA FISH imaging

Cells were imaged with a Nikon Ti Eclipse epifluorescnet microscope equipped with perfect focus (Nikon), a 100x VC DIC lens (Nikon), fluorescent filters for YFP, DAPI, TMR and CY5 (Semrock), an X-cite 120 fluorescent light source (Excelitas), and an Orca Flash 4v2 CMOS camera (Hamamatsu). The microscope was controled by Micro-Manager program [3].

#### 1.3 Image aquisition for time-lapse and single molecule RNA FISH imaging

For live cell time-lapse microscopy the microscope was operated in constant focus mode. Bright field images were taken every 10 s with an exposure time of 10 ms and the YFP fluorescent images are taken every 1 minute with an exposure time of 20 ms. For single molecule RNA-FISH microscopy, z-stacks of images from fixed yeast cells for bright field, DAPI, TMR and CY5 were taken with each image in the z-stack separated by 200 nm. For each sample, muliple positions on the slide were imaged to ensure large numbers of cells.

#### 1.4 Sample preparation for live cell time-lapse microscopy

1.5 ml of yeast cells with Hog1-YFP in log-phase growth (OD=0.5) were pelleted by centrifugation, resuspended in 20 *μ*l CSM medium, and loaded into a flow chamber as described in [4]. The media passing through the flow chamber was removed through a syringe (New Era Pump Systems) connected to a pump (pump rate 0.1 ml/minute) at the exit of the flow chamber. On the input of the flow chamber, a two-way valve was connected to switch between a beaker with CSM media and a beaker with CSM medium with a fixed concentration of NaCl (0.2M or 0.4M).

#### 1.5 Image analysis for time-lapse microscopy

Each image from the fluorescent tagged Hog1-kinase (Hog1-YFP) was used to generate a background image by running a disk smoothing filter over the image. The background image was subtracted from the orginial YFP image to enhance contrast. This image was then smoothed twice with a median filter. From this image the mean pixel intensity for pixels above 50 counts was computed. This value serves as a threshold to convert the YFP image into a binary image in which high intensity YFP signal that is enriched in the nucleus, is used as a nuclear marker. The bright field image was smoothed with a disk filter to generate a background image. The background image was subtracted from the orginial brightfield image to enhance contrast. The processed YFP image and the processed brightfield image were combined using morphological reconstruction, and cells were segmented with a watershed algorithm. Cells that are too small, too big, or too close to the image borders were removed. This process was repeated for each image. After segmentation, the centroid of each cell was computed and stored. Next, the distance between each centroid for each of the two consecutive images was compared. The cells in the two images that have the smallest distance were considered the same cell at two different time points. This whole procedure was repeated for each image resulting in single cell trajectories. For each cell the average per pixel fluorescent intensity of the whole cell (Iw) and of the top 100 brightest fluorescent pixels (It) was recorded as a function of time. In addition, fluorescent signal per pixel of the camera background (Ib) was reported. The Hog1 nuclear enrichment was then calculated as Hog1(t) = [(It(t) - Ib) / (Iw(t) - Ib)]. The single cell traces were smoothed and subtracted by the Hog1(t) signal on the beginning of the experiment. Next, each single cell trace was inspected and cells exhibiting large fluctuations due to poor cell segmentation and cell tracking were removed. During the time points when no fluorescent images were taken, the fluorescent signal from the previous time point was used to segment the cell. Taking images every 10 s with an exposure time of 10 ms using bright field imaging did not result in photobleaching of the fluorescent signal and resulted in better tracking reliability because cells had not moved significantly since the previous image. For cell volume measurements, each time trace was removed of outlier points that resulted from segmentation uncertainties. Volume change relative to the volume at the beginning of the experiment was calculated to compare cells of different volumes. For both, the single-cell volume and Hog1(t) fluorescent traces, the median and the average median distance (standard deviation of the median) was computed to put less weight on sporadic outliers due to the image segmentation process. The final time-lapse microscopy data set consist of 246 (0.2M NaCl) and 167 (0.4M NaCl) cells. Each time course was measured in duplicates or triplicates.

#### 1.6 Sample preparation for single-molecule RNA-FISH

Yeast cell culture (*BY4741* WT) in log-phase growth (OD = 0.5) was concentrated 10X (OD = 5) by a glass filter system with a 0.45 μm filter (Millipore). Cells were exposed to a final osmolyte concentration of 0.2 M and 0.4 M NaCl, and then fixed in 4 % formaldehyde at time points 0, 1, 2, 4, 6, 8, 10, 15, 20, 25, 30, 35, 40, 45, 50, 55 minutes. At each time point, cells (5 ml) in the corresponding beaker were poured into a 15 ml falcon tube containing formaldehyde, resulting in cell fixation. Cells were fixed at +20C for 30 minutes, then transferred to +4C and fixed overnight on a shaker. After fixation, cells are centrifuged at 500 xg for 5 minutes, and then the liquid phase was discarded. The cell pellet was resuspended in 5 ml ice-cold Buffer B (1.2 M sorbitol, 0.1 M potassium phosphate dibasic, pH 7.5) and centrifuged again. After discarding the liquid phase, yeast cells were resuspended in 1 ml Buffer B, and transferred to 1.5 ml centrifugation tubes. Cells were then centrifuged again at 500 xg for 5 minutes and the pellet was resuspended in 0.5 ml Spheroplasting Buffer (Buffer B, 0.2 % βmercaptoethanol, 10 mM Vanadyl-ribonucleoside complex). The OD of each sample was measured and the total cell number for each sample was equalized. 10 μl of 2.5 mg/ml Zymolyase (US Biological) was added to each sample on a +4C block. Cells were incubated on a rotor for 20-40 minutes at +30C until the cell wall was digested. The cells turn from an opaque to a black color if they are digested. Cells were monitored under the microscope every 10 minutes after addition of Zymolyase and when 90 % of the cells turned black, cells were transferred to the +4C block to stop Zymolyase activity. Cells were centrifuged for 5 minutes, then the cell pellet was resuspended with 1 ml ice-cold Buffer B and spun down for 5 minutes at 500 xg. After discarding liquid phase, the pellet was gently re-suspended with 1 ml of 70 % ethanol and kept at +4C for at least 12 hours or stored until needed at +4C. Hybridization and RNA-FISH probe preparation conditions were applied as published [5]. Table S1 contains the RNA-FISH probes for *STL1* and *CTT1*. Each time course was measured in duplicate or triplicate.

#### 1.7 Image analysis for single-molecule RNA-FISH

To segment cells and to define the cells nuclear and cytoplasmic compartment the following steps were taken. For each DAPI image stack, a maximum intensity projection was generated. A DAPI intensity threshold was picked by eye to identify the maximum number of nuclei in the image. Based on this threshold, the image was then converted into a binary image. Connected regions (nuclei) that were too big or too small were removed. Connected regions in the image were then labeled by individual numbers resulting in individual numbered nuclei. Next, for each connected region three individual DAPI thresholds were computed as 40 (low threshold) and 50 (middle threshold) % of the difference between the maximum DAPI signal minus the background DAPI signal. Through this iterative and cell specific thresholding, cell-to-cell differences in nuclei DNA concentration and DAPI staining were taken into account. For the transcription site analysis, the low threshold was used. For the determination of the nuclear and cytoplasmic joint probability mRNA distribution, the middle threshold was used. For each cell, the entire DAPI image stack was converted into a binary image stack using the cell specific DAPI thresholds resulting in 3D thresholded nucleus and cytoplasm. After the nucleus had been segmented, the last 5 images of the bright field image stack were maximum projected to generate the cell outlines. A background image was generated by running a disk smoothing filter over this image. The background image was subtracted from the maximum projected image to enhance contrast. This image was then combined with the thresholded DAPI image using morphological reconstruction and cells were segmented with a watershed algorithm. Cells that were too small, too big, or too close to the image borders were removed.

To identify RNA spots, fluorescent thresholds for RNA-FISH images were determined for each dye based on a single image plane, for each position imaged and for each sample. For a given dye, the mean threshold value was calculated based on the thresholds identified from each image in the data sets. This threshold was very reproducible from image to image and from person to person. After the threshold had been determined for each dye, each image in the stack was Gaussian filtered to remove noise, and then filtered with a laplacian of a Gaussian filter to detect punctuate in the image. These filtered images were then converted into binary images using the previously determined threshold. For each pixilated signal, the regional maxima was determined, which identifies the xyz-position of the RNA spot within each 3D image stack. This processes was repeated for each of the TMR and CY5 image stacks. The number of RNA spots for each cell in the cytoplasm or the nucleus was determined by applying the mask of segmented nucleus and cytoplasm on the filtered RNA-FISH image stacks in 3D. The numbers of RNA molecules in the nucleus and cytoplasm were counted for each individual cell.

In order to determine the number of transcripts of each transcription site, we determined the total intensity of each RNA spot in the cell. We also fitted each RNA spot with a 2D Gaussian function to determine the RNA spot intensity and found very good agreement between the fitted and the total RNA spot intensities. The total RNA-FISH data set consists of a total of 65454 single cells (25511 at 0.2M NaCl and 39943 at 0.4M NaCl).

#### 1.8 RNA-FISH data analysis

For each time point, a distribution of mRNA molecules in single cells was determined as the marginal distribution of total *STL1* and *CTT1* mRNA or as the joint probability distribution of nuclear and cytoplasmic RNA. The distributions were also summarized in a binary data set of ON and OFF cells. Cells with three or more mRNA molecules were considered ON-cells for quantification of the ON-fraction. The population average was computed as the mean marginal cytoplasmic or nuclear RNA for each mRNA species.

To quantify transcription site intensities, we recorded the intensity of all spots in the nuclei for all cells. The brightest nuclear spots of each cell were labeled as potential transcription sites, and were temporarily removed from the data set. We then computed the median intensity for the remaining nuclear spots, and we used this median value to define the equivalent mature nuclear mRNA fluorescence intensity. Next, we re-examined the brightest RNA spot in each nuclei (i.e., the potential transcription sites), and we computed the fluorescence intensity in units of mature mRNA intensities. Potential transcription sites that had intensities greater than the equivalent of two mature mRNA molecules were labeled as active transcription sites. This approach was applied to all cells from a single experiment. From this analysis, we computed the fraction of cells with an active transcription site as a function of time, the single-cell distributions of full length RNA transcripts per transcription site as a function of time, and the average number of full length RNA transcripts per transcription site as a function of time.

### 2 Methods–Computational

All computational analyses have been performed using Mathworks MATLAB. All analysis codes are downloadable at XXX and required data sets are available at YYY^1^. Once the files have been downloaded, all results and Figures (2-4 and S1-S11) can be generated by (1) opening Matlab and navigating to the main directory, and running the function Munsky_2017_Main.M. This function will open a simple interface that allows the user to generate all figure and to manipulate system parameters.

#### 2.1 Parameterizing the Hog1p signal

To model the temporally-varying Hog1p localization signal, we adopt the empirical model from [5]. In this model, the level for Hog1p(t) enrichment is assumed to have the form:

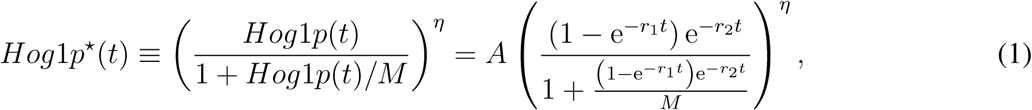

where *A* and *M* define the saturation height and midpoint and are the same for all salt levels. See the supplemental information of [5] for the derivation of this function.

The parameters {*r*_1_, *α*, *η*, *A*, *M*} have been fit to match the newly measured Hog1p nuclear enrichment levels as functions of time, and osmolyte concentrations. The parameters are given in Table S2, and the corresponding fits to the new experimental data are shown in Fig. S1. This time-varying signal has been used as an input to the gene regulation models below.

#### 2.2 Gene regulation model

To capture the spatial stochastic expression of *STL1* or *CTT1* mRNA, we extend the four-state *Hoglp*-activated gene expression model identified in [5] to account for spatial localization of mRNA in the nucleus or cytoplasm (see Fig. 1A, 1B). The model consists of a single gene that can occupy one of four different transcriptional states (*S*_1_, *S*_2_, *S*_3_, *S*_4_) and two chemical species: nuclear mRNA (*m*_nuc_), and cytoplasmic mRNA (*m*_cyt_). The gene can transition between the four possible states (*S_j_* → *S_k_*) with the rates, {*k_jk_*}. The rate of the state transition *S*_2_ → *S*_1_ depends on the level of Hog1p according to the simple saturated linear form,

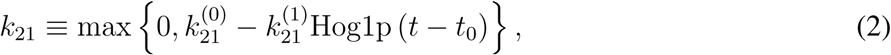

where *t* is the time following the addition of NaCl, and *t*_0_ is a time delay required to capture the short period between transcription initiation and dispersion of mature mRNA^2^. All other state transition rates are constant in time. Each state *S_j_* allows transcription of mRNA in the nucleus with rate, *k_i_j__*. For the spatial model, nuclear mRNA are transported to the cytoplasm with rate, *η*_r_. mRNA are assumed to degrade at the rate γ. For the spatial model, a second degradation rate γ_nuc_ is assumed for the nuclear region.

In total, there are 13 non-spatial or 15 spatial parameters in the model:

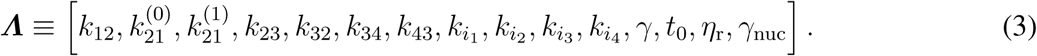

For parameter uncertainty quantification, we assume an independent, log-uniform prior on all parameters, and all parameter searches are conducted in logarithmic space.

#### 2.3 Computation of statistical moments of gene regulation fluctuations

In this section, we provide the sets of ordinary differential equations (ODE’s) that describe the moments of the proposed model.

##### 2.3.1 First and second moments

The analysis of the first two moments (i.e., the means, variances and covariances) are a special case of the general form for the higher order moments described below in Section 2.3.2. Let x(*t*) denote the state of the system at any given time:

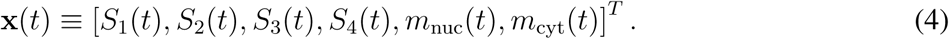

For this study, x is a vector of five (non-spatial) or six (spatial) discrete, non-negative integers. Chemical reactions are random events that take the system from one state to another: x_*i*_ → x_*j*_. Each *μ*^th^ reaction can be described by its stoichiometry vector, *s_μ_*, which describes the state change that occurs for that reaction (i.e., ***s****_μ_* = x_*j*_ – x_*i*_ if the *μ*^th^ reaction goes from x_*i*_ to x_*j*_) and its propensity function *w_μ_*(x, *t*)*dt*, which describes the probability that such a reaction would occur in the infinitesimally small time step, *dt*.

For the *non-spatial model*, there are six unique state transition stoichiometries, one transcription reaction, and one degradation reaction, for a total of eight unique stoichiometry vectors. These can be combined into the stoichiometry matrix:

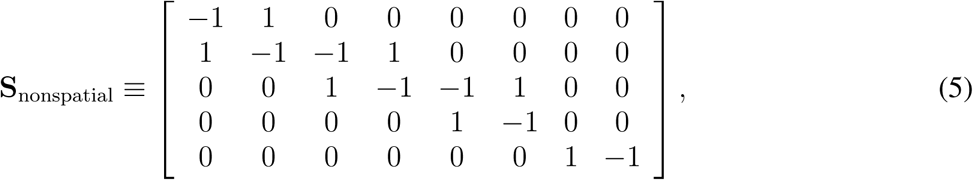

in which each row corresponds to one of the state variables, and each column corresponds to a reaction. The corresponding propensity functions can also be written in matrix-vector form for the *nonspatial models* as:

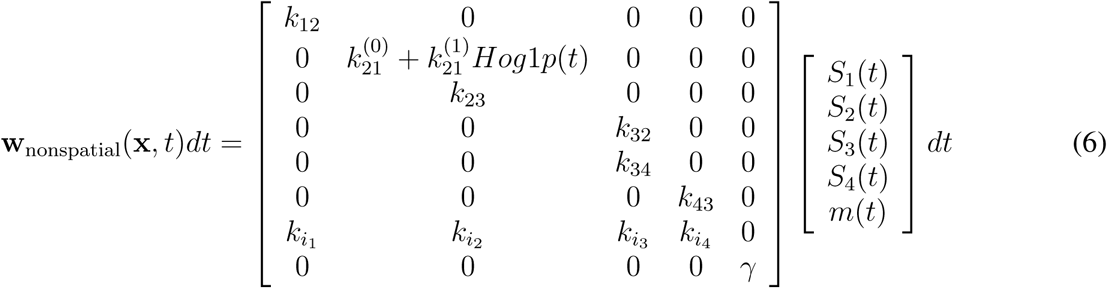

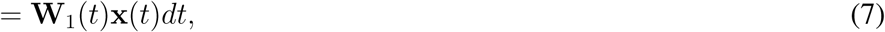

Similarly, the spatial model has an additional two reactions corresponding to transport from the nucleus to the cytoplasm and an extra degradation term for the nuclear mRNA. The stoichiometry matrix for the *spatial model* can be written:

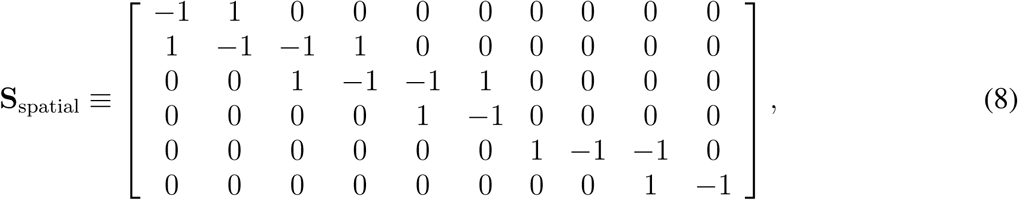

and the corresponding propensity functions can also be written as:

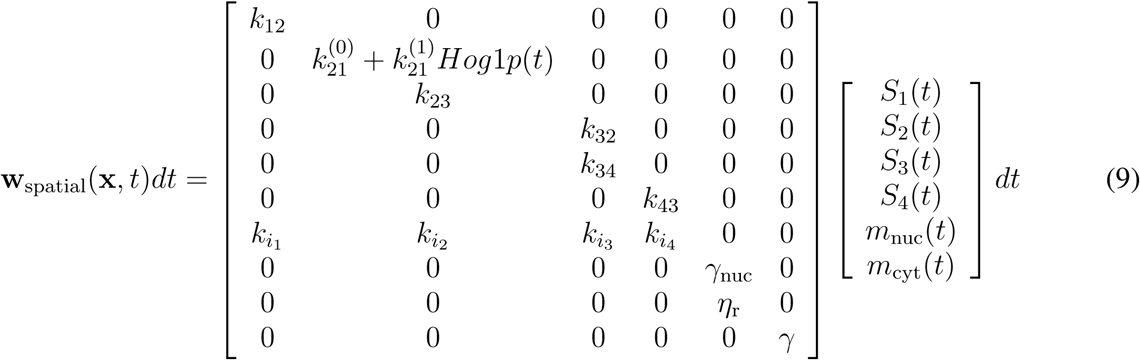

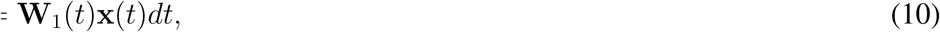

With these definitions, the time-varying means ***μ***(*t*) = 𝔼{x(*t*)} of all species and the corresponding covariance matrix ***Σ***(*t*) = 𝔼{[x(*t*) – ***μ***(*t*)][x(*t*) – ***μ***(*t*)]^*T*^} can be computed by integrating the linear ordinary differential equations:

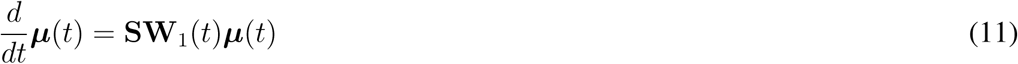

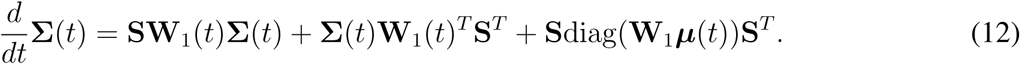

This integration has been performed using Mathworks Matlab.

##### 2.3.2 Higher moments of gene regulation fluctuations

In order to compute the likelihood to observe the first two moments of gene regulation fluctuations, it is necessary to compute the third and fourth moments of the fluctuations as well. These higher moments can also be described by a set of linear ODEs as follows. Let 𝔼{x^*β*^} denote a higher order uncentered moment of the random vector x:

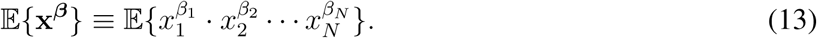

With this definition the dynamics of 𝔼{x^*β*^} can be shown to evolve according to the differential equation [6,7]:

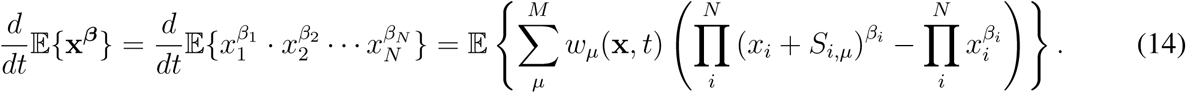

In the special case where all propensity functions are first order, then Eqn. 14 represents a closed system of equations, which can be integrated to solve for the first four uncentered moments.

#### 2.4 FSP computation of probability densities of gene regulation fluctuations

To compute the full time varying probability distributions, we use a slightly different notation. Let y(*t*) = [*i*, *j*, *k*] denote the state of the system where *i* is the index of the current gene state, *S_i_*, *j* is the number of nuclear mRNA, and *k* is the number of cytoplasmic mRNA. Let *P*(y(*t*)|***Λ***, **P**_0_) denote the probability of the state y at time *t* conditioned upon the model ***Λ*** and the initial probability distribution at *t* = 0, given by the vector **P**_0_. For the 4-state model examined in this study, the index *i* can take four different values, *i* ∈ {1, 2, 3, 4}, and *m*_nuc_ and *m*_cyt_ could be any non-negative integer value, {0,1, 2,…}. The chemical master equation [8] can be written:

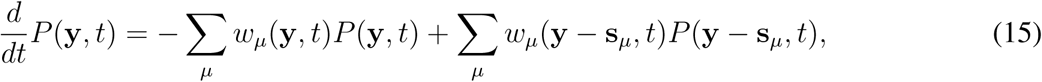

where the sum is over all possible reactions.

We enumerate all possible states in the set {y_1_, y_2_,…}, and we define the corresponding probability density mass vector as: **P**(*t*) = [*P*_1_(*t*), *P*_2_(*t*)…]^T^. Using the Finite State Projection analysis [9,10], we truncate the full probability mass vector to a finite set where the number of mRNA in the nucleus and cytoplasm are each bounded by finite numbers, *N*_nuc_ and *N*_cyt_. We utilize use a nested enumeration, given by *y*_*I*(*i,j,k*)_ = [*i*, *j*, *k*], where *I*(*i*, *j*, *k*) = *N*_states_(*N*_nuc_ + 1)*k* + *N*_states_*j* + *i*.

With this enumeration, the Chemical Master Equation can be written in vector form as: 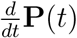 = **Q**(*t*)**P**(*t*). We solve this equation in Matlab using the stiff ODE solver, ode15s. For comparison of the model to the experimental mRNA distributions, it is necessary to convert the **P**(*t*) into a marginal distribution of nuclear and cytoplasmic mRNA numbers. This is relatively easy to do, since

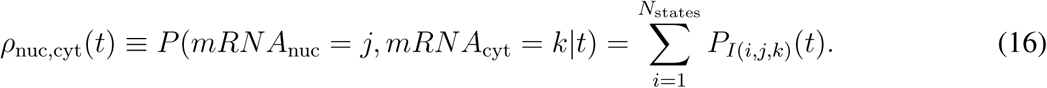

The FSP approach and moments analyses were verified to match all moments up to fourth order, each within 0.002%.

Now with this definition, we can compare model predictions and experimental data as described below.

#### 2.5 Computation of Likelihood Functions – Moments Analyses

For a population sample of *N*_S_ cells in which the mRNA has been measured as {*x*_1_, *x*_2_,… *x*_N_s__}, the measured sample mean (*μ_S_*) and unbiased sample variance estimate 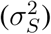 are:

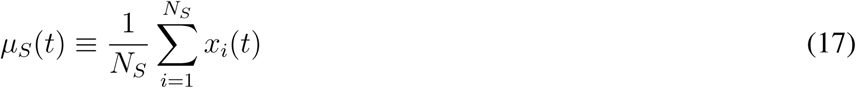

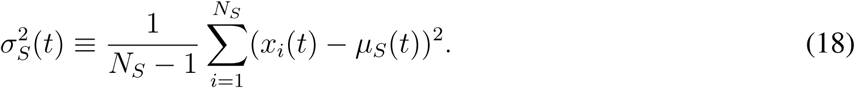

##### 2.5.1 Application of Central Limit Theory

The moment analyses discussed in this article (i.e., Eqns. 12 and 14) provide exact expressions for the dynamics of the model’s means, variances, covariances and higher moments. Because the reaction rates are all linear, these moment equations are closed, and the moments can be computed with no assumption on the distribution shape. To compare these computed moments to measured experimental data requires computation of the likelihood that the measured moments (e.g., means or covariance matrix) would have been observed by chance given that the model were correct. The standard practice for such a computation is either to assume Gaussian deviations or to apply the Central Limit Theorem (CLT) as follows. Let *x̃*_*i*_ be a set of independent and identically distributed random variables with finite mean, *μ*, and finite variance, σ^2^, and let *ỹ* be the average of *N_s_* such random numbers.

According to the CLT, as *N_s_* becomes very large, the distribution of *ỹ_Ns_* will approach a normal distribution with a mean of *μ* and a variance of σ^2^/*N_S_*. The CLT allows us to estimate the likelihood of the observed moments as discussed in the following sections. We note that if *x̃_i_* is normally distributed, then *ỹ_N_S__* would be normally distributed for any number of cells. However, in the case where *x̃_i_* is very far from normally distributed (e.g., when its distribution has a very long asymmetric tail) then a larger number of cells will be required before the CLT becomes valid. Such is the situation observed for the later time points in Fig. 3, which result in poor estimation of the moments-based likelihood function.

##### 2.5.2 Likelihood to observe measured means

###### 2.5.2.1 Univariate Case – Non-Spatial Model

Assuming a finite variance, σ^2^(*t*), and a large numbers of cells, *N_S_*, the CLT states that the sample mean of a population of cells will have a distribution that can be approximated by a normal distribution with mean *μ_m_* and variance σ^2^(*t*)/*N_S_*:

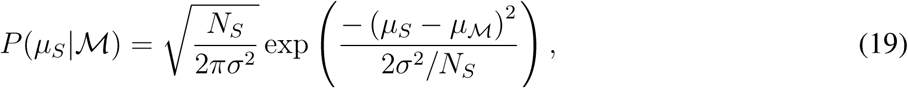

where 𝓜 denotes the model under consideration. Taking the logarithm of this likelihood function yields:

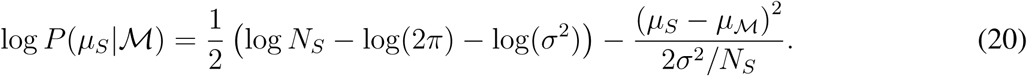

We can now compute the log likelihood of the averaged data given our model (up to a constant, *C*_1_), provided that: (1) we can compute the mean, and (2) the variance is known:

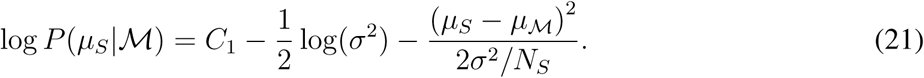

In the case where we are only using the average measurements to constrain the model, we replace the variance with the sample variance estimate:

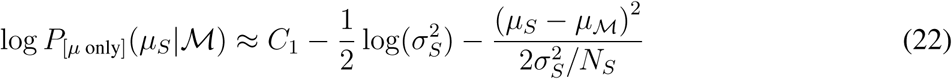

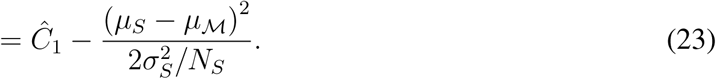

In this expression, all terms that do not depend upon the model mean, *μ_𝒨_*, have been lumped into the constant *Ĉ*_1_.

###### 2.5.2.2 Multivariate Case – Spatial Model

The spatial model has *n* = 2 observables: nuclear and cytoplasmic mRNA levels, but the analysis is quite similar. When the number of cells in each sample, *N_S_*, is large, the distribution of the sample mean vector, ***μ**_S_*, given the model can be approximated by a multivariate normal distribution with mean, ***μ**_𝒨_*, and covariance (1/*N_S_*)**Σ**. The likelihood of observing the measured mean at a given time point can be written

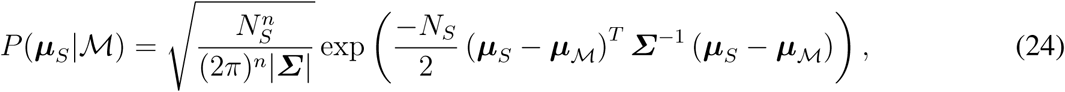

where the notation |***Σ***| denotes the determinant of the covariance matrix, ***Σ***. Taking the logarithm of this likelihood and collecting terms that do not depend on ***μ**_𝒨_* or ***Σ*** into a constant yields:

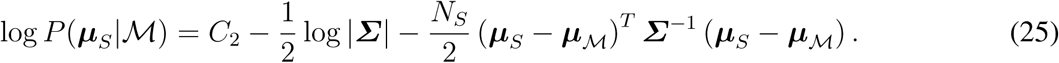

To attempt to identify the model without computation of the second moments, we must approximate *Σ* with the measured sample covariance estimate:

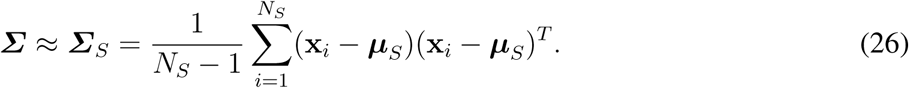

For this special case, the likelihood reduces to:

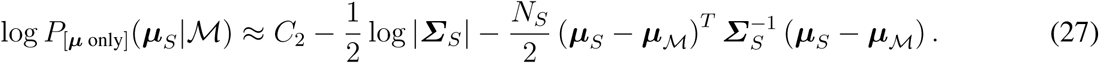

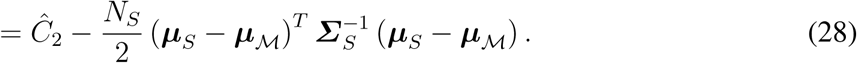

As in the previous case, all terms that do not depend upon the model mean, ***μ**_𝒨_*, have been lumped into the constant *Ĉ*_2_.

##### 2.5.3 Likelihood to observe measured (co)variances

To estimate model parameters using the gene expression variance, it is necessary to compute the likelihood that we would observe the measured *sample variance* given the model. For a single observable variable (e.g., non-spatial model for a single mRNA species), the sample variance, 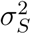, will be a random variable that depends upon the distribution of the number of mRNA per cell at the corresponding time-point.

In this work, we consider two different CLT-based approaches to approximate the likelihood to observe the measured sample variances. The first method assumes that the likelihood of the observed sample variance can be well approximated by a 𝒳-squared distributed random variable, which is independent of the sample mean. This approximation would be exact if the underlying distribution had a Gaussian distribution. This approximation of the likelihood depends only upon the model’s computation of the first two moments. The second method approximates the sample mean and sample (co)variances as multivariate Gaussian random variables [7], whose covariance depends upon the first four moments of the analysis (thus the need for the extended moments analysis described in Section 2.3.2). These analyses are described in greater detail as follows:

###### 2.5.3.1 Univariate Case - 𝒳^2^ distribution approximation

Using the assumption that the sample population is sufficiently large that 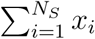 has a normal distribution, one could approximate the probability of observing the measured sample variance using the 𝒳-squared distribution with κ = *N_s_* - 1 degrees of freedom:

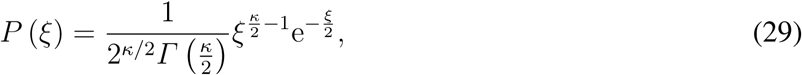

Where 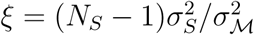, and κ = *N_s_* – 1. Or by changes of variables, this can be written:

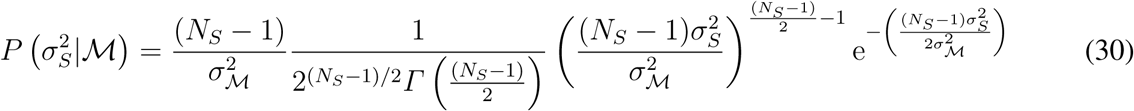

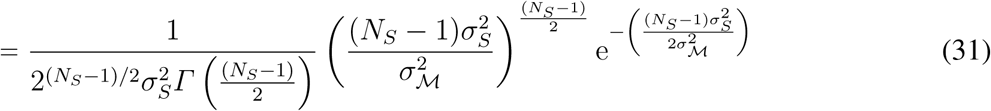

The logarithm of the likelihood of the observed sample variance 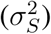 given the model variance (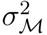)

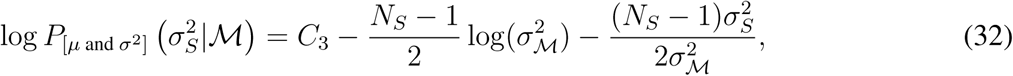

where *C*_3_ is the collection of all terms that do not depend upon 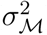.

###### 2.5.3.2 Multivariate Case – Wishart distribution approximation

To estimate the likelihood function for the observed variance and covariances in the multivariate case, we use a generalization of the 𝒳-squared distribution (Eqn. 31), known as the Wishart distribution [11]. To define this likelihood function, let ***Σ**_𝒨_* represent the covariance matrix generated by the model. Let **X**_*s*_ represent an *N_s_* by *n centered matrix* of the measured data for *N_s_* different independent measurements of n distinct chemical species:

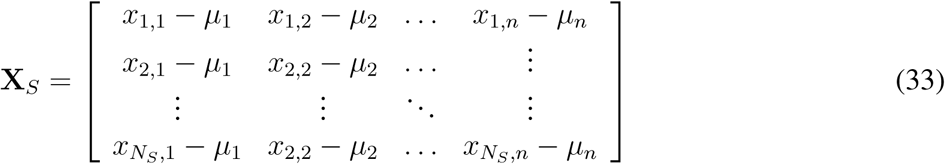

Define S_*s*_ as the *n* × *n scatter matrix* of the data, 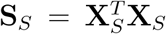. With these definitions, the Wishart distribution over the range of possible S can be written:

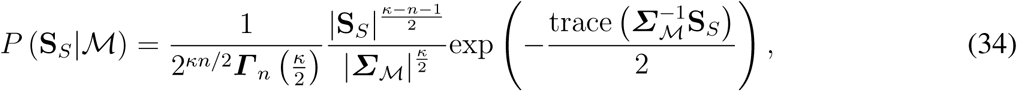

where κ = *N_s_* – 1 is the degrees of freedom.

Taking the logarithm of Eqn. 34 and collecting all terms that are independent of ***Σ***_𝒨_ yields:

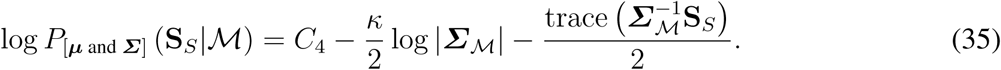

Using the relationship ***Σ**_s_* = (*N_s_* – 1)^−1^S_*S*_, this reduces to a form analagous to that for the univariate case in Equation 32:

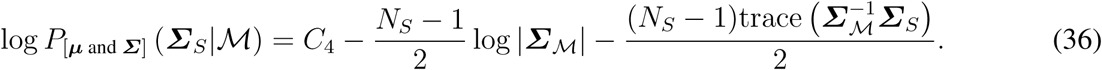

###### 2.5.3.3 Likelihood to observe measured means and (co)variances using the 𝒳-squared/Wishart approximation

Assuming a (multivariate) normal distribution for the number of nuclear and cytoplasmic mRNA, the sample means and the sample variances (covariance matrix) would be statistically independent. Therefore, the log-likelihood to match both statistics for the univariate case is the sum of Eqns. 21 and 32 over all time points *k*; = {1, 2, …, *K*}:

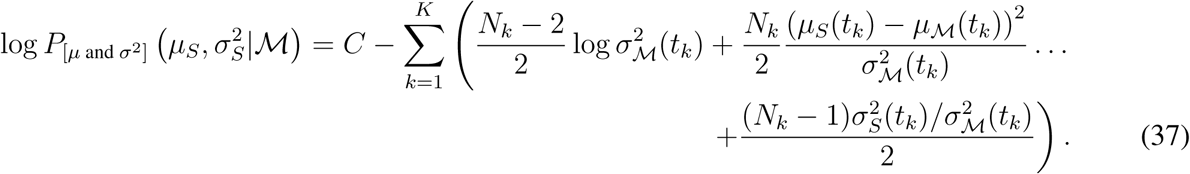

Similarly, in the multivariate case, the log-likelihood to match both the measured sample mean vector, *μ*_S_, and the measured sample covariance matrix, *Σ*_S_, can be found from the sum of Eqns. 28 and 36:

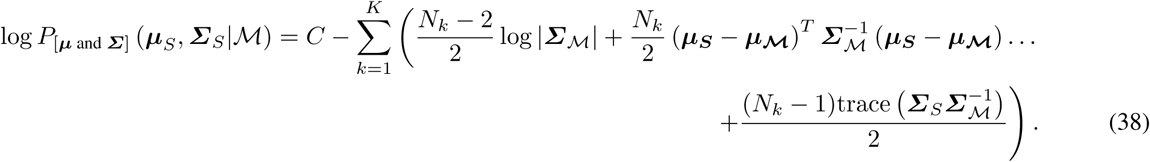

Naturally, Eqn. 38 reduces to Eqn. 37 in the case of a single chemical species.

###### 2.5.3.4 Likelihood to observe measured means and (co)variances using a multivariate Gaussian approximation

For strongly skewed distributions (e.g., those measured in this study), the sample mean and sample variance both depend strongly upon the experimental sampling of the distribution tails. As a result, these statistics are not statistically independent. To account for this interdependence, the CLT may be used to approximate the joint likelihood of the sample means and sample (co)variances as a multivariate normally-distributed random vector. For two measured species (e.g., nuclear and cytoplasmic mRNA), this vector has five elements (z ≡ [*μ*_1_, *μ*_2_, *σ*_11_, *σ*_12_, *σ*_22_]), and its probability distribution can be written in the form [7]:

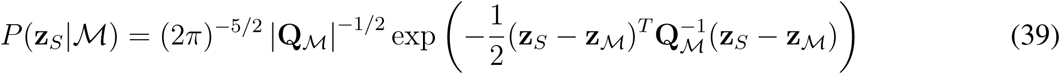

where z_*S*_ is the vector of measured means, variances and covariances; z_𝒨_ is the corresponding model-predicted average of those statistics; and Q_𝒨_ is the model-predicted covariance matrix for those statistics. The matrix Q_𝒨_ can be defined in terms of the second through fourth order moments as follows [7]:

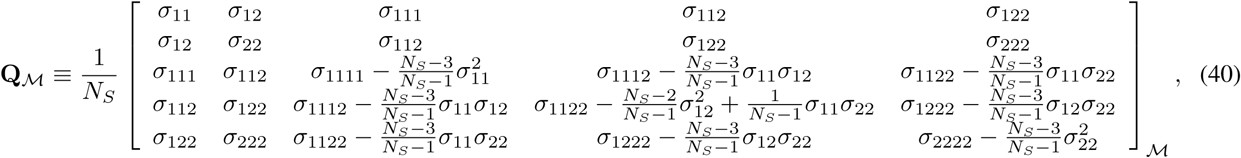

where σ_*β*_1__…*β_N_* is used to denote the (∑*β_i_*) centered moment:

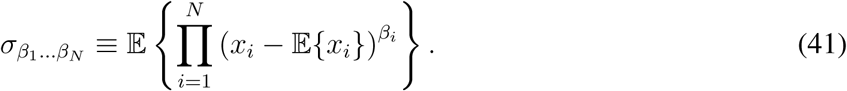

Under this approximation, the logarithm of the the likelihood can be written:

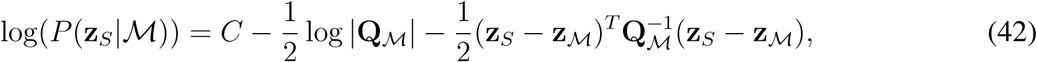

and the log-likelihood for all time points, *k* = 1,…, *K* can be written:

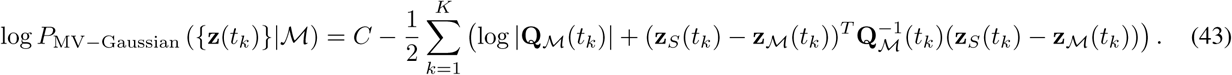

##### 2.5.4. Comparison of the 𝒳^2^/Wishart distributions and multivariate Gaussian approaches for the likelihoods of the first two moments

For the analysis of *CTT1* mRNA, the extension of the moments analyses to include the third and fourth moments led to identification of parameters that were 10^93,900^ times less likely to account for the full measured mRNA distributions compared to the analysis using the FSP analysis (Table S5 and Section 2.8). Moreover, this result was 10^63,300^ times less likely to account for the distribution data than was the model identified by the much simpler 𝒳^2^ approach (Table S5). Similar deleterious effects were observed for the spatial *STL1* and CTT1 analyses, although inclusion of the third and fourth moments led to some improvement in the case of non-spatial *STL1* analysis, for which the skewness had the greatest effect as shown in Figs. 3 and S7.

Recall from the main text that the model is unable to fit the means and variances exactly due to the difficulty to estimate those statistics due to high skewness. In order to compensate for this mismatch, the extended first-four moments approach identifies very large 3rd and 4th moments. However, these higher moments are not directly constrained by the data, and they are severely over-estimated (as opposed to under estimated as they were in the 𝒳^2^ and Wishart distribution based approaches). As a result, the extended model provided excelent fits to the means, but fits to all other statistics are much worse as shown in Fig. S6. In principle, this could be corrected by using data from the 3rd and 4th moments to add additional constraints to the model, but such an approach would be problematic for two reasons: (1) the measurement of the 3rd and 4th moments are prone to even greater sampling error than are the first two moments; and (2) in order to compute the likelihood to observe the third and fourth moments requires computation of even higher moments, which increases the computational complexity and makes such analysis impractical for the current model.

Because the extended moments analysis requires greater computational effort to compute while providing worse predictions to the full distributions in most cases, we have restricted the majority of our discussion to the simpler moments-based analyses.

#### 2.6 Likelihood to observe measured distributions

Because the FSP approach provides a direct computation of the model’s probability distribution, the FSP-based computation of the likelihood of the measured data is much more straightforward, and it does not require any assumptions of the distributions’ shapes. To define this likelihood function, suppose that *n_c_* cells, *c* = [1, 2,…, *n_c_*], were measured for a given experiment and time point, and each cell was found to have exactly *m*_nuc_(*c*) copies nuclear mRNA and *m*_cyt_(*c*) copies cytoplasmic mRNA. Suppose that a model with parameter set *Λ*, predicts that for each time and experiment, the probability that a given cell has exactly *m*_nuc_(*c*) nuclear mRNAs and *m*_cyt_(*c*) in the corresponding conditions is *p*(*m*_nuc_(*c*), *m*_cyt_(*c*) |*Λ*). The total likelihood of all observations, *L*(**D**|**Λ**), is the product over every cell, or

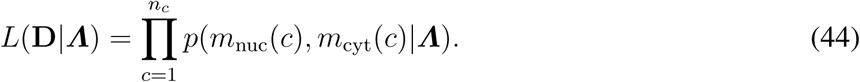

Now that we know how likely it is that the data comes from a model and a given set of parameters, ***Λ***, our goal is to find the parameter set, ***Λ***_Fit_, which maximizes this likelihood (or equivalently the logarithm of this likelihood):

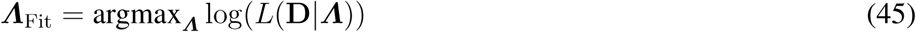

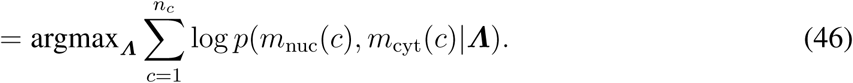

Under the assumption that cells are independent, and because each cell is measured only once using the smFISH technique, the log-likelihood that the model matches all experiments at all time points is the sum of the log-likelihoods of the individual experiments and time points.

#### 2.7 Parameter searches to maximize likelihood

In order to conduct parameter searches, we use iterative combinations of simplex based searches (e.g., Matlab’s “fminsearch” function) and genetic algorithm based searches. All parameters were defined to have positive values, and searches were conducted in logarithmic space. The searches are run multiple times from different starting parameter guesses leading to many tens of millions of function evaluations (> 4 × 10^7^ evaluations of the means and moments analyses, > 5 × 10^6^ evaluations of the non-spatial FSP distributions, and > 1 × 10^6^ evaluations for the more computationally expensive analyses of extended moments and the spatial FSP distributions). Parallel fits were conducted on clusters of more than 128 processors at a time allowing for the consideration of several millions of model/ parameter/ experiment combinations per day.

#### 2.8 Comparison of results for different likelihood functions (Table S5)

Table S5 shows the relative log-likelihood values for different likelihoods that compare the means, means and variances, extended moments, or full distributions. Each row corresponds to a different combination of gene and identification strategy. Each column corresponds to a different likelihood function (i.e., means, means and (co)variances, extended moments, or full distributions). Values presented are log_10_ of the actual likelihoods relative to the best value found for that specific objective function.

The ‘Distributions’ *column* of Table S5 quantifies the relative log-likelihood that the identified parameter set matches *all data* of that type (i.e., spatial or nonspatial). More negative numbers denote worse matches to the full data. For example, in the non-spatial analysis of *CTT1*, the 𝒳^2^ and extended moments-based approaches were respectively 10^30,600^ and 10^93,900^ times less likely to account for the full data than was the model identified by the FSP approach. In other words, the full data was 10^63,300^ times less likely to have come from the model identified using the extended moments than it was to have come from the simpler 𝒳^2^-based analysis.

The Full FSP distributions *row* of Table S5 shows the relative likelihood of the Full FSP Distribution parameters when compared to the best fit for each of the different likelihood functions. This quantity provides an estimate of the probability that each likelihood function would have discovered the FSP parameters by chance. One interesting observation from this perspective is that although the extended moments analyses produce worse fits to the full data, this effect is somewhat mitigated by the fact that the extended analysis yields greater uncertainties (see also Figs. 4B, S8-S10). In other words, the extended analysis leads to worse fits to the full data, but this is coupled with lower confidence in these poor results.

#### 2.9 Metropolis Hastings algorithm to quantify parameter uncertainties

To quantify the parameter uncertainties, we used the Metropolis Hastings algorithm [12], which is a standard Markov Chain Monte Carlo (MCMC) algorithm for parameter uncertainty estimation. All MCMC parameter explorations were conducted in logarithmic parameter space, and all analyses (i.e., means, means and variances, or distributions, both spatial and non-spatial) used the same proposal distribution for the MCMC chains. This proposal distribution was a symmetric normal distribution (in logarithmic space) with a variance of 0.005 (also in logarithmic space). For each parameter proposal, a random selection of parameters were selected to change, where each parameter had a 50% chance to be perturbed. The first half of each chain was discarded as an MCMC burnin period, and all chains were thinned by 90%.

##### 2.9.1 Convergence

MCMC chains for the means, simpler moments analyses, and non-spatial FSP analyses were run for MH chains with combined lengths of >25,000,000 parameter evaluations. MCMC analyses for the more computationally expensive extended moment analyses were run for combined lengths of >1,500,000 parameter evaluations. MCMC analyses for the most computationally expensive spatial distribution analyses were run for a combined length of >3,700,000 parameter evaluations. To evaluate convergence of the MCMC analyses, we have split the multiple MCMC chains into two disjoint sets. To illustrate the achieved convergence, Figures S8 shows the distributions of likelihood values for these two independent MCMC compilations for each analysis and for *STL1* and *CTT1.* Thus, all 32 MHA analyses (2 replicas × 2 genes × 4 statistical analyses × 2 spatial analyses) were confirmed to have sampled similar distributions of log-likelihood space.

##### 2.9.2 MCMC Results

Figures S9 and S10 summarize the biases and pairwise parameter uncertainties for the *STL1* and *CTT1* analyses and for each of the eight different analyses (means, means and variances, extended moments, and distributions) × (spatial and non-spatial).

The vectors in Figs. S9 and S10 represent the fold over- or under-estimation bias for the corresponding parameter. The matrices in Figs. S9 and S10 represent the variances (diagonal entries) and covariances (off-diagonal entries) of the parameter estimation uncertainty, as estimated using the Metropolis Hastings algorithm. For example, the figure shows that non-spatial means analysis overestimates the *k_i_3__* and γ parameters (‘+’ symbols in the corresponding biases), is highly uncertain in the estimate of *k_i_3__* (yellow box in the covariance matrix diagonal corresponding to that parameter), and the uncertainties in parameters *k_i_3__* and γ are positively correlated (‘+’ symbol in corresponding off-diagonal entries of the covariance matrix). The open red ellipse plotted in Figs. 4A visually depicts the same information by plotting the 90% confidence parameter interval, assuming a lognormal posterior distribution with the MCMC-estimated pairwise covariance matrix.

The *total bias* shown in Figs. 4C and S11C was defined by the euclidean distance in logarithmic space according to the expression:

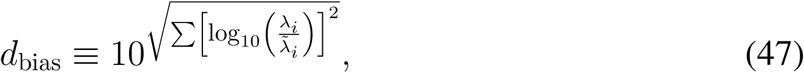

where {λ_*i*_} is the average of the *i*^th^ estimated parameter, and 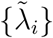 is the corresponding ‘true’ parameters (i.e., the maximum likelihood estimate using all data and the spatial FSP analysis). The summation is taken over all parameters *except* 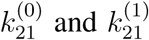, which provide a non-unique determination of the Hog1p saturation effect, *k*_*i*_1__, which is nearly zero and therefore highly uncertain in logarithmic space, and *t*_0_ whose interpretation is different for the spatial and non-spatial models. Conceptually, the value *d*_bias_ provides a quantification of the overall ± fold changes for the collective parameters compared to their ‘true’ values.

The *total uncertainty* shown in Figs. 4B and S11B was defined by the trace of the covariance matrix. As was the case for the total bias quantification, the trace is taken over all parameters *except* 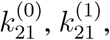 *k*_*i*1_, and *t*_0_.

##### 2.10 Fisher Information analysis

The Fisher Information Matrix (FIM), *I*(***Λ***), and its inverse, the Cramér-Rao bound, is an analysis technique that describes the lower bound on the variance of an unbiased estimator. Recent works [7,13,14] have applied this tool in a biological context, looking at model sensitivity, robustness, identifiability, and experiment design. Here, we are concerned with the ability to identify models, which (1) only consider the means or the means and variances, and (2) assume a Gaussian likelihood function for the sample means and variances. The FIM is defined for the parameter vector ***Λ*** [λ_1_, λ_2_,…, λ_*p*_] and likelihood *L*(**D**|***Λ***) as

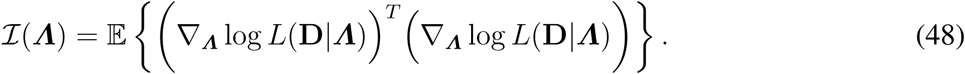

In this work, we have compared parameter uncertainties using posterior distributions of the parameters. The Cramér-Rao inequality states that the variances of these distributions must be less than or equal to the inverse of the FIM, *I*(***Λ***)^−1^. A model is unidentifiable if *I*(***Λ***)^−1^ does not exist, i.e. if the FIM is singular [15].

For the models that consider means ***μ***, means and variances, ***μ*** and *Σ,* we have assumed Gaussian forms of the likelihood function, Eqns. 28 and 38. Under this assumption, the FIM is well-known. For a model which only describes the mean of the distribution, it is given by

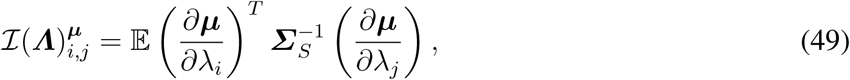

and when both means and variances are to be estimated with the model, the FIM is

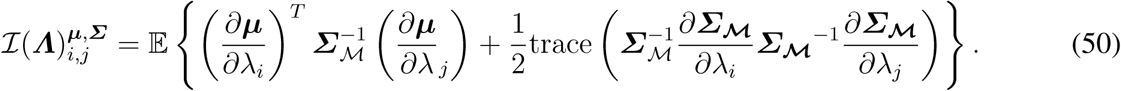

Given a set of independent samples at each experimentally measured time point, the FIM can be constructed at each time point by integrating Eqns. 11 and 12, and the total information is the sum over all the time points. For models of *μ* and *μ* and *Σ*, we computed the FIM and found that them to be invertible, indicating that the parameters are in principle able to be identified.

##### 2.11 Predictions of Transcription Site activity

We developed two analyses to predict transcription site (TS) activity: a simplified theoretical analysis of average active TS activity and an extended FSP analysis of distributions of polymerases on a given TS. These are described below. For both the data and the model, TS sites were considered to be ON if their predicted or measured intensities were greater than twice the intensity of a single mature mRNA.

###### 2.11.1 Simplified theoretical model of average TS polymerase loading

Using the mRNA distribution analyses described above, we identified transcription initiation rates, {*k*__*i*1__, *k*_*i*_2__, *k*_*i*_3__, *k*_*i*_4__} for the *CTT1* and *STL1* mRNA. For an elongation rate of *k*_elong_ and an mRNA length of *L*, each polymerase takes a time of τ_elong_ = *L/k*_elong_ to complete transcription. Thus, polymerases that initiate transcription in the time window (*t* – τ_elong_, *t*) will be present at the TS at time *t*. For each gene, we selected the fastest identified transcription rate *k*_i–max_ = max{*k*_*i*_1__, *k*_*i*_2__, *k*_*i*_3__, *k*_*i*_4__}. Assuming that an *active* TS is in the state with the maximum transcription rate, the average number of polymerases per active TS, 〈*n*_pol_〉, can be computed as:

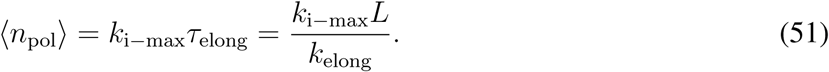

In practice, L was chosen as the length from the first smFISH probe on the 5’ end of the mRNA to the transcription termination site. In light of the fact that the smFISH probes are nearly equally distributed along the mRNA, we assume that nascent mRNA are on average half the length of their mature counterparts. Under this assumption we related *n*_pol_ = 2*n*_nascent_, which allows us to relate the elongation rate to the observed spot intensities (in terms of mature mRNA) as:

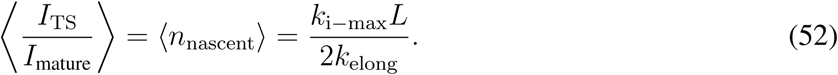

###### 2.11.2 Extended FSP model of nascent transcription

Next, we extended Finite State Projection approach in a form similar to that in [16]. To compute the distribution for the number of polymerases at the TS, we first compute the distribution of the gene states at the earlier time, *t* – τ_elong_. Using this distribution of gene states at *t* – τ_elong_ and ignoring previous initiation events and excluding degradation of partially described mRNA, we use the FSP analysis derived above (but with no degradation or transport) to solve for the distribution of elongating polymerases per TS at the time *t*. We assume that each partially transcribed mRNA is at a random location in the gene, and it therefore has an effective intensity uniformly distributed between zero and the equivalent of one mRNA. To find the distribution of TS spot intensities with *N*_poly_ polymerases, we take the convolution of *N*_poly_ independent random variables, each with a uniform distribution between zero and one.

To confirm the FSP analysis, we also simulated the identified models using an adapted form of the Stochastic Simulation Algorithm (SSA, [17]) but with a modification similar to that proven in [18]. In each run of the SSA, we recorded the times of every simulated transcription initiation event. With this simulated information and an assumption of deterministic mRNA elongation with a given elongation rate, we compute the distance traveled by each polymerase as a function of time. If that length is longer than the length of the gene, then that polymerase is assumed to have completed transcription, in which case the mRNA is assumed to have dissociated from the TS. Otherwise, the length of the elongating mRNA and the known smFISH probe placements (Table S1) are used to determine how many probes are located along the partially transcribed nascent mRNA for that particular polymerase. The total TS intensity is then computed as the sum of the intensities for all polymerases on the gene in that cell and at that point in time.

The standard SSA approach [17] assumes constant propensity functions. In order to allow for time varying rates in the propensity functions (i.e., Eqn. 2), we added a fast reaction to the system. The stoichiometry of this reaction was zero, such that each firing of this null reaction does not affect the state of the system [18], but it does allow for frequent updates to the propensity functions. The rate of this reaction was set to 1*s*^−1^, which is more than two orders of magnitude faster than the characteristic rates of the Hog1p fluctuations in Eqn. 1. Figure S12 shows excellent agreement between the FSP and SSA analysis of the TS activity for *STL1* and *CTT1* transcription under 0.2M and 0.4M NaCl osmotic shock conditions.

###### 2.11.3 Identification of mRNA elongation rate

In order to identify the transcription elongation rate, we computed the TS intensity distribution for *CTT1* at each point in time for 0.2M and 0.4M NaCl osmotic shock using the previously identified parameters (Table S4) and one free constant to describe the average elongation rate, *k*_elong_. We then computed the probability that the observed distributions of *CTT1* TS intensities could have originated from this model at all time points and conditions. We then maximized this likelihood with respect to the rate kelong for the different experimental replicas and NaCl concentrations to determine the uncertainty in this parameter. Using the simplified theoretical model, which does not account for transitions between active and inactive periods, we found an *upper bound* on the *CTT1* elongation rates to be 91±9Nt/s. Using the more detailed spatial FSP approach, we found the *CTT1* elongation rates to be 63±14Nt/s. For both cases, the uncertainty is given as the standard error of the mean using the five experimental replicas (two for 0.2M NaCl and three for 0.4M NaCl). We then fixed the elongation rate to be 63Nt/s, and we used this rate in conjunction with the previously identified parameters to predict the TS intensity distributions for *CTT1* and *STL1* as functions of time in both osmotic shock conditions (Figs. 4D and S11D-H).

**Figure S1:**
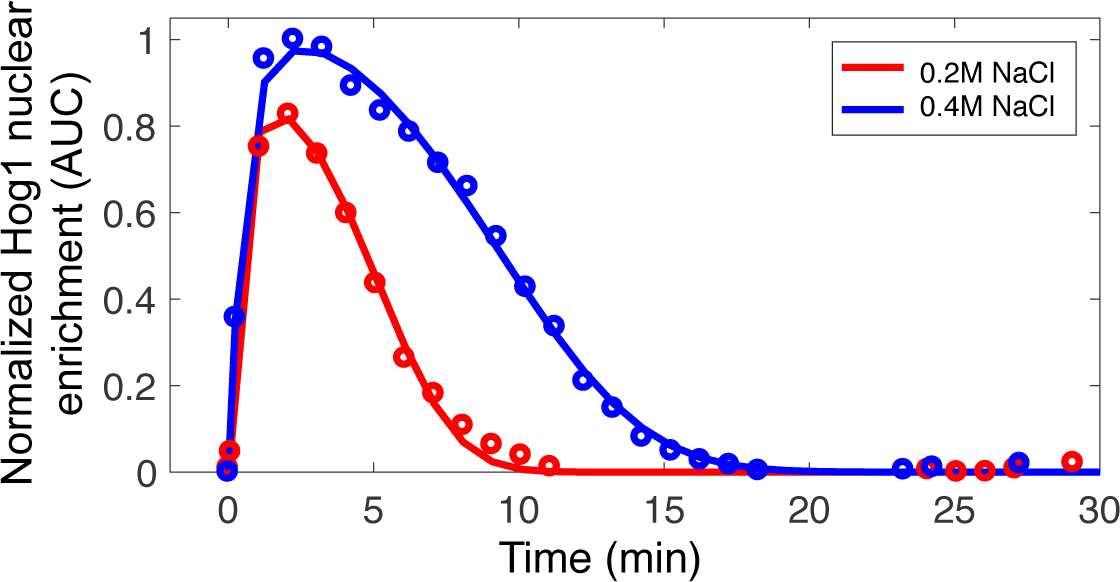
Normalized Hog1 nuclear enrichment after activation of the HOG-pathway with different NaCl concentrations. Solid lines denote the model (Eqns. 1) fit to the data points (symbols) for the two different osmolyte concentrations.

**Figure S2:**
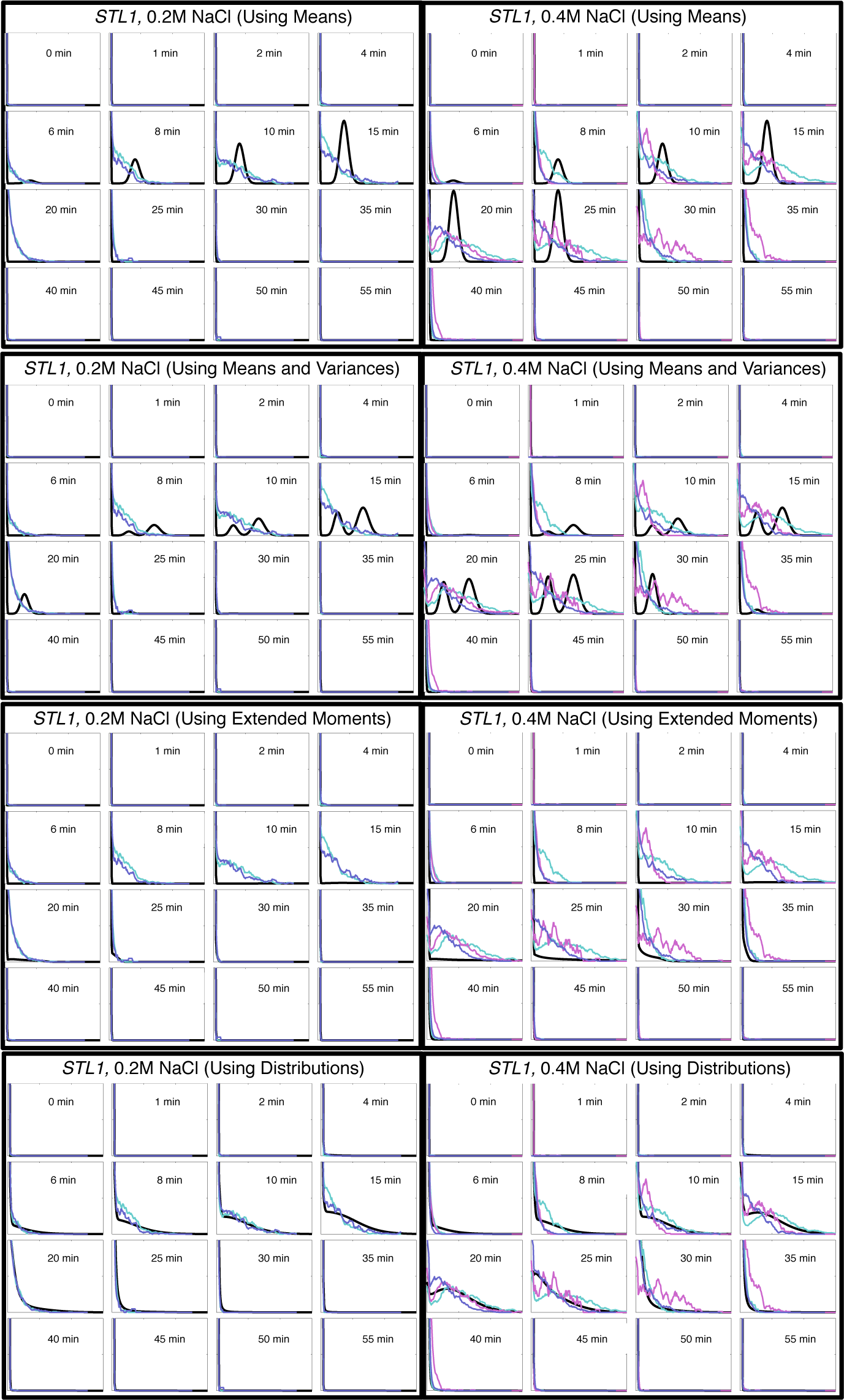
Experimental and model-generated distributions for the number of *STL1* per cell versus time for 0.2M NaCl (left) and 0.4M NaCl (right) osmotic shocks. Distributions have been generated using the models identified using means only (top row), means and variances using the 𝒳^2^ approach (second row), means and variances using the extended moments analysis (third row) or full distributions (bottom row). Model results are shown in black. Experimental data replicates shown in colors. Limits of all x-axes are 0 to 150 copies of mRNA. Limits of all y-axes are probability mass of 0 to 0.04.

**Figure S3:**
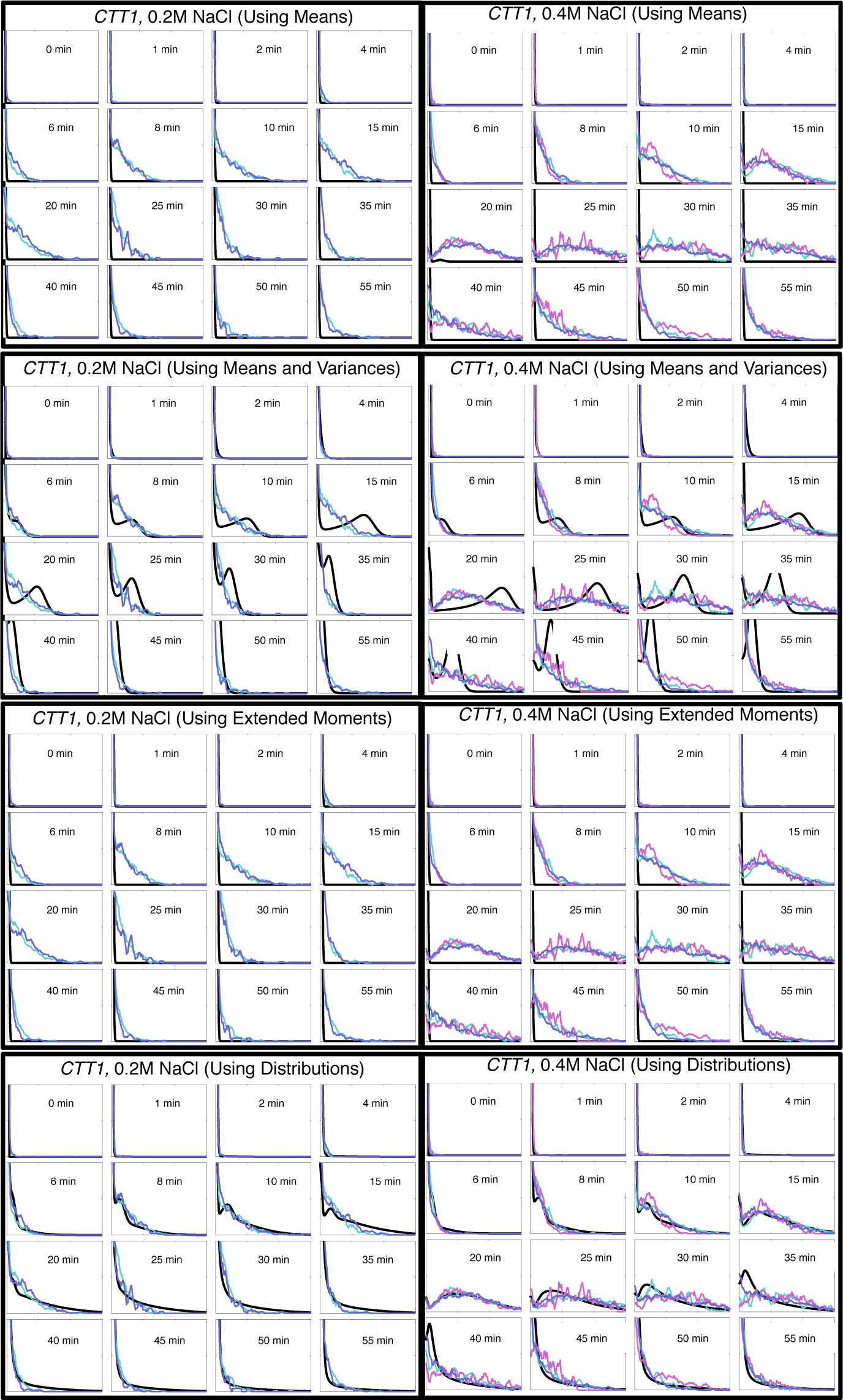
Experimental and model-generated distributions for the number of *CTT1* per cell versus time for 0.2M NaCl (left) and 0.4M NaCl (right) osmotic shocks. Distributions have been generated using the models identified using means only (top row), means and variances using the 𝒳^2^ approach (second row), means and variances using the extended moments analysis (third row) or full distributions (bottom row). Model results are shown in black. Experimental data replicates shown in colors. Limits of all x-axes are 0 to 150 copies of mRNA. Limits of all y-axes are probability mass of 0 to 0.04.

**Figure S4:**
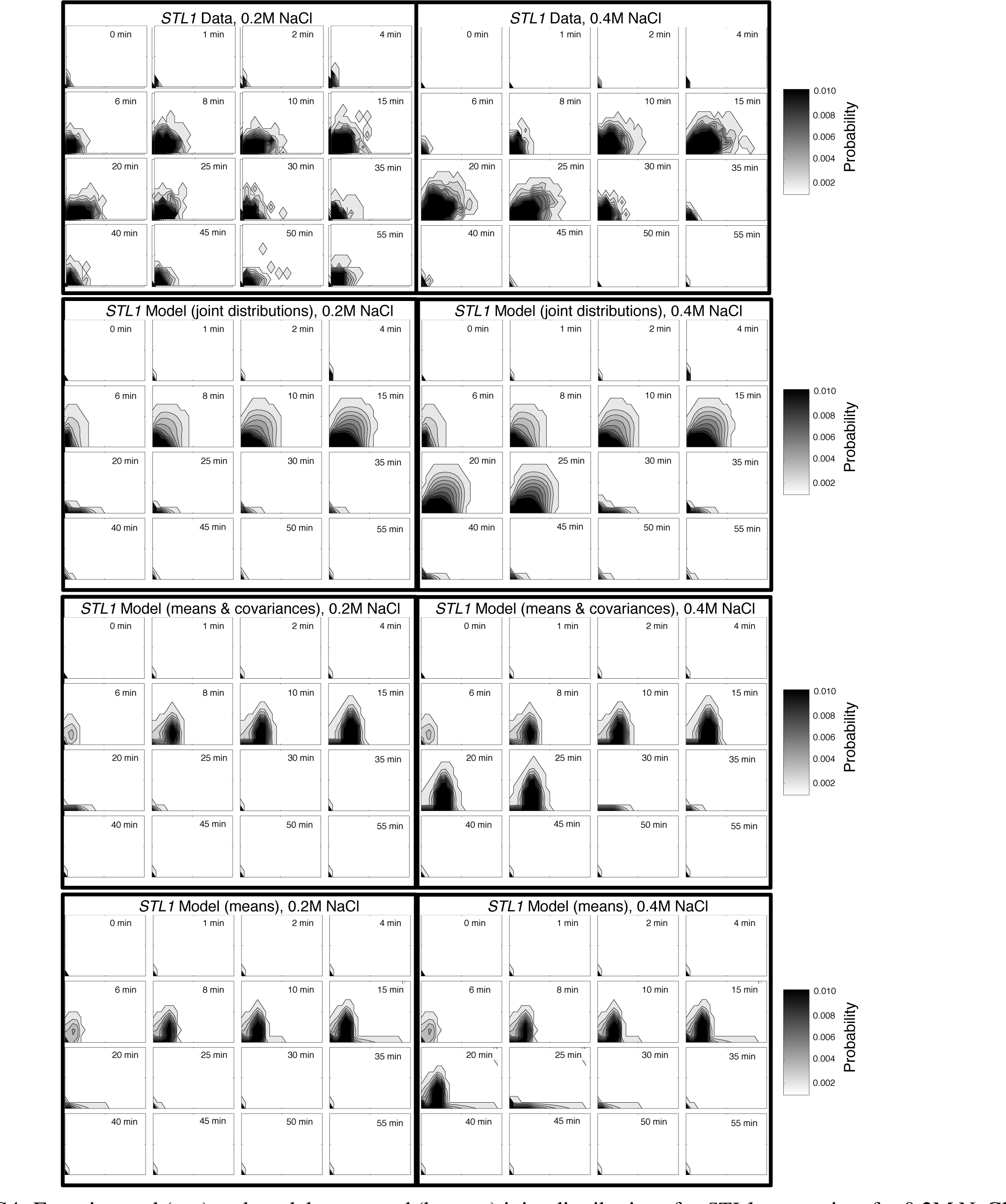
Experimental (top) and model-generated (bottom) joint distributions for *STL1* versus time for 0.2M NaCl (left) and 0.4M NaCl (right) osmotic shocks. Distributions have been generated using the model identified from the full joint distributions (row 2), the means and covariances (row 3) or the joint means (row 4). Limits of all x-axes are 0 to 200 copies of cytoplasmic mRNA. Limits of all y-axes are 0 to 10 copies of nuclear mRNA. All plots use the same contour levels (see legend to right). See Table S3 for the corresponding parameter sets.

**Figure S5:**
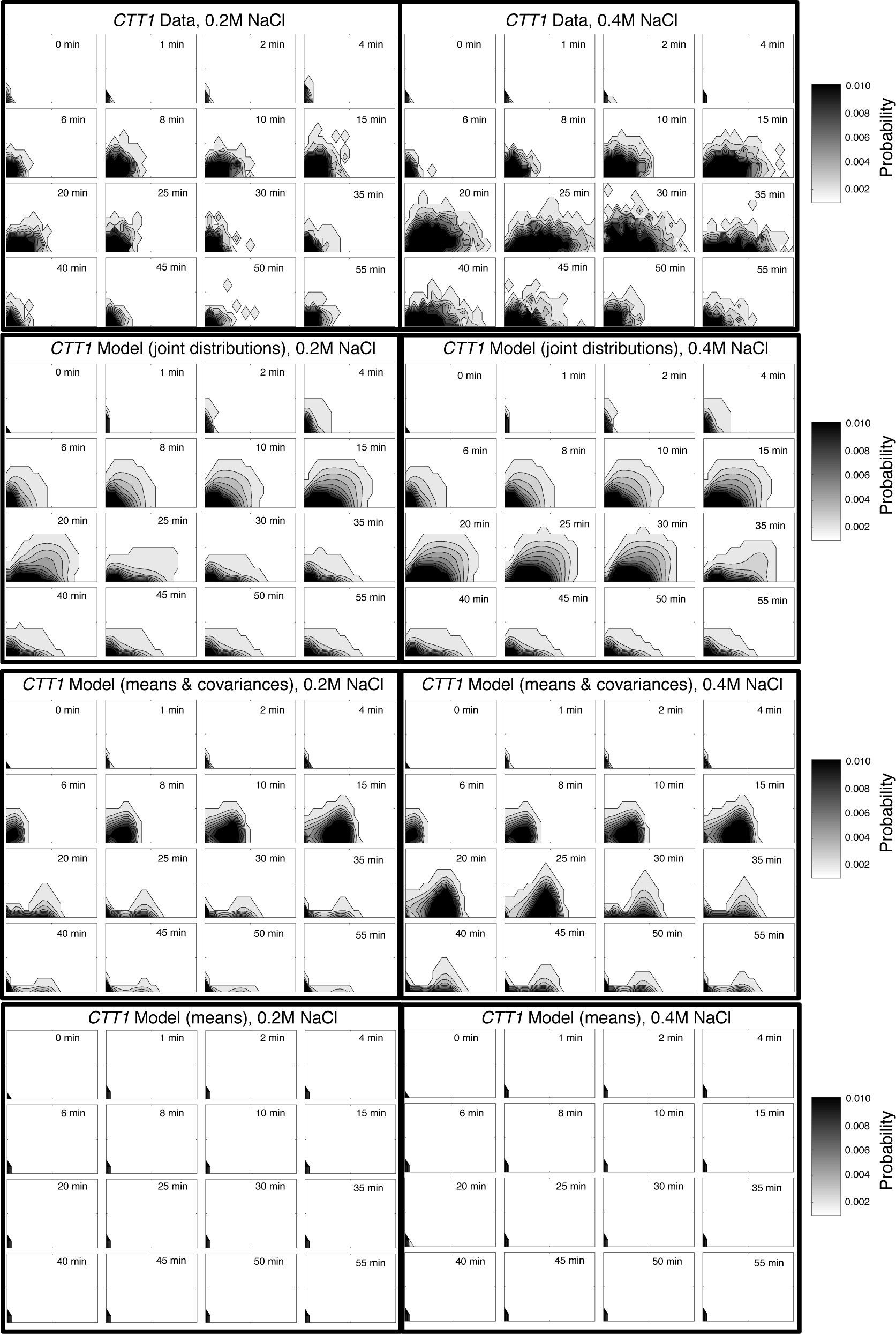
Experimental (row 1) and model-generated (rows 2-4) joint distributions for *CTT1* versus time for 0.2M NaCl (left) and 0.4M NaCl (right) osmotic shocks. Distributions have been generated using the model identified from the full joint distributions (row 2), the means and covariances (row 3) or the joint means (row 4). Limits of all x-axes are 0 to 200 copies of cytoplasmic mRNA. Limits of all y-axes are 0 to 10 copies of nuclear mRNA. All plots use the same contour levels (see legend to right). See Table S4 for the corresponding parameter sets.

**Figure S6:**
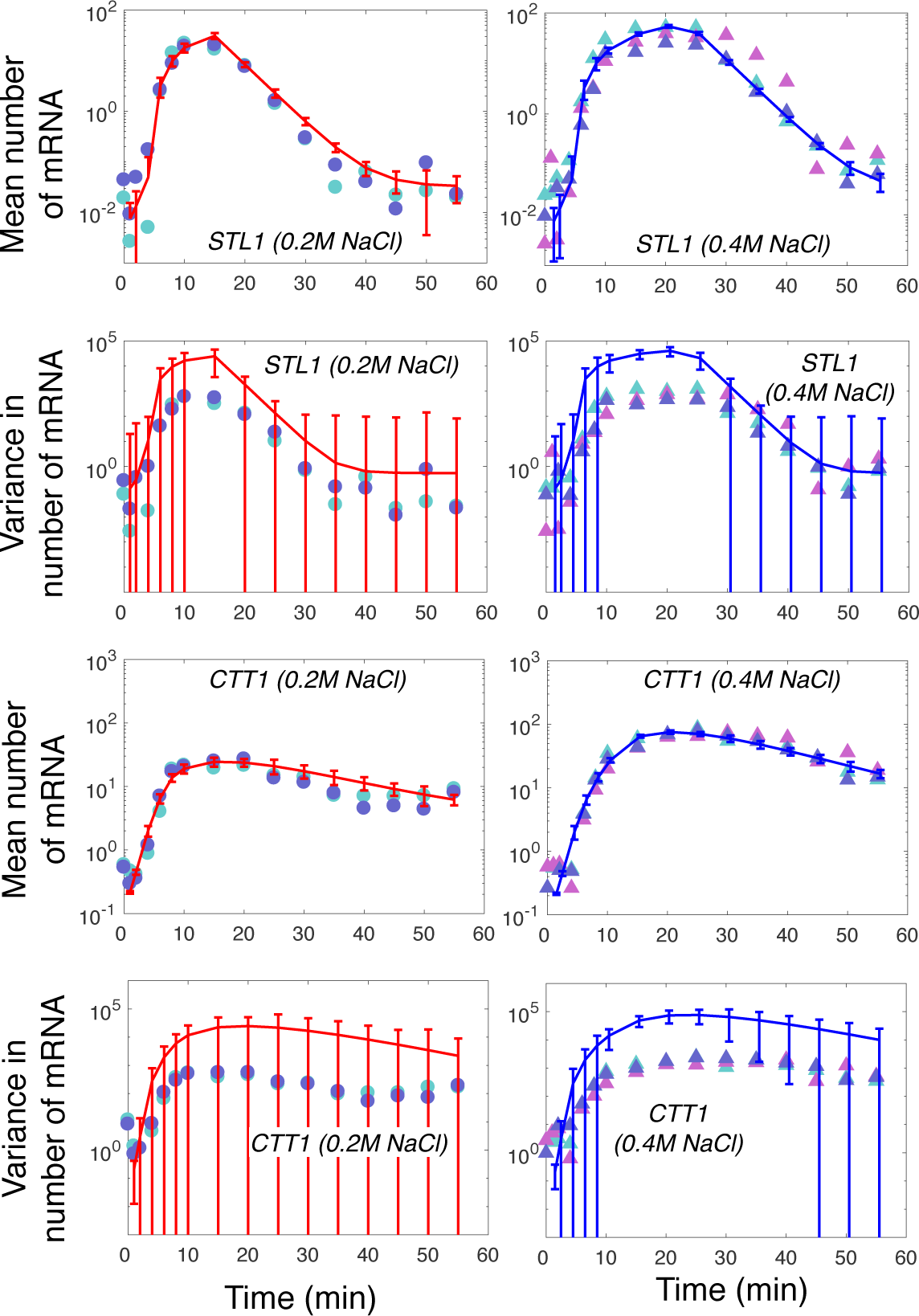
Best fit to the means and variances using the extended moments analysis for *STL1* and *CTT1* for 0.2M NaCl (left, 2 biological replica) and 0.4M NaCl (right, three biological replica) osmotic shock. Error bars denote the expected errors of one standard error above and below the estimated value for the corresponding statistic. According to the likelihood function defined in Section 2.5.3.4, the standard error of the sample mean is given by 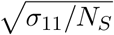, and the standard error of the sample variance is given by 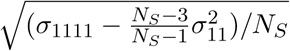, where *N_s_* is the sample size corresponding to the total measured number of cells for that time point and condition). Experimental data replicates are shown by the circles and triangles.

**Figure S7:**
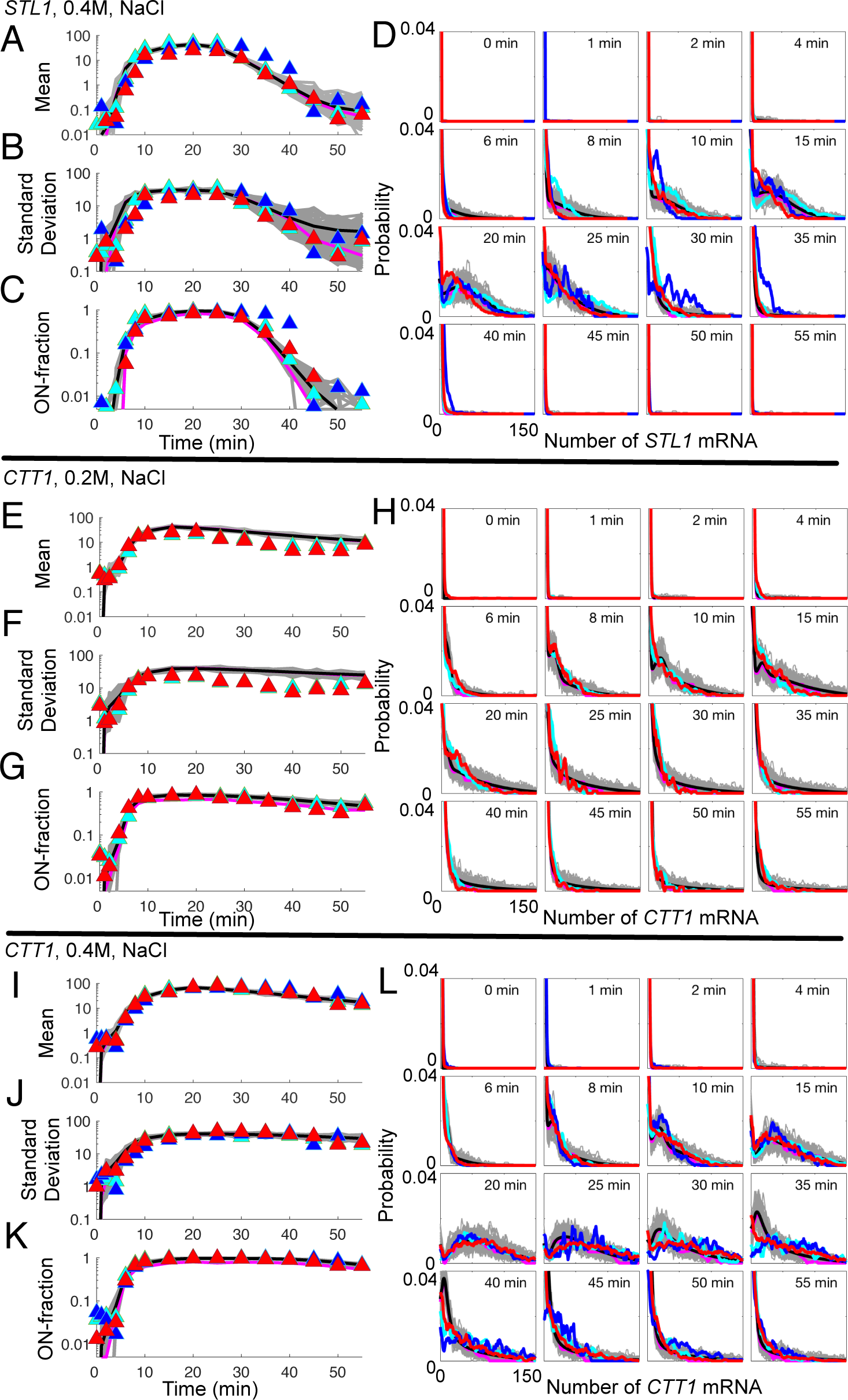
Means (A,E,I), standard deviations (B, F, J), ON-fractions (C, G, K) and probability distributions (D, H, L) for *STL1* at 0.4M NaCl osmotic shock (A-D), *CTT1* at 0.2M NaCl osmotic shock (E-H), and *CTT1* at 0.4M NaCl osmotic shock (I-L). In all panels, theoretical values from the fitted model are in black, representative simulated samples of 200 cells apiece are in gray, median statistics of the simulated samples are in magenta; and experimental data are in red and cyan (and blue for 0.4M NaCl). See also Figure 3 in the main text.

**Figure S8:**
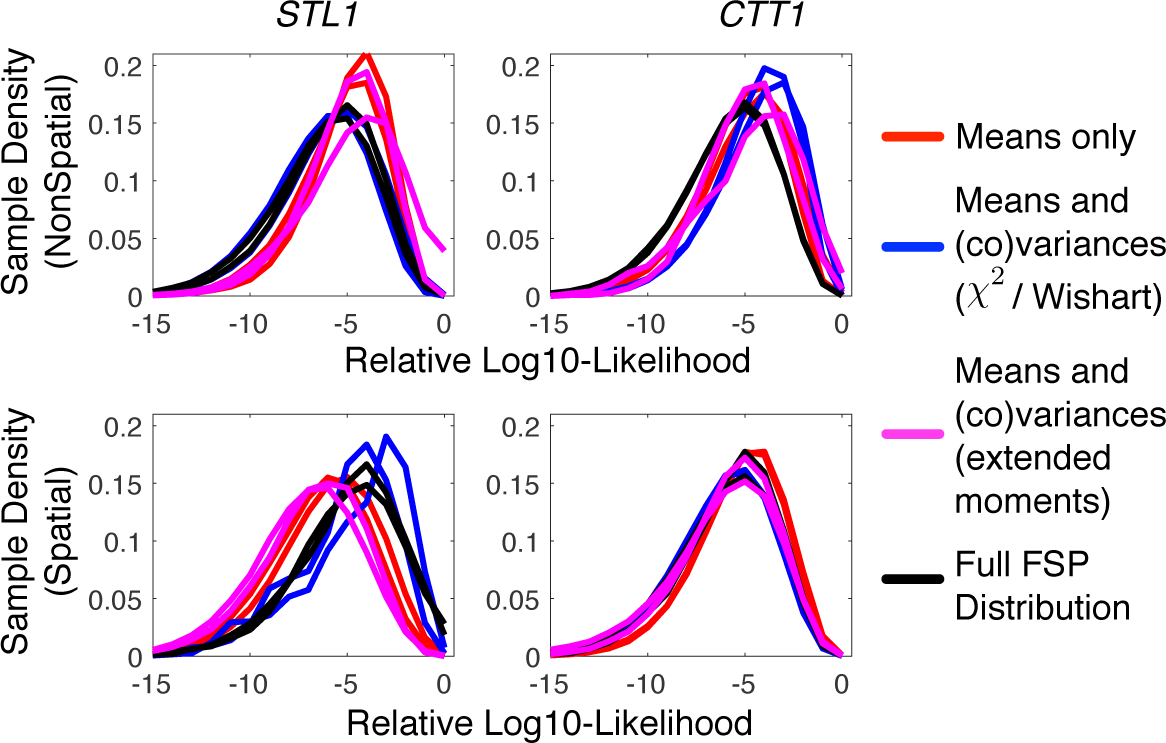
Distributions of log_10_-likelihoods that have been sampled by the Metropolis Hastings analysis for *STL1* (left) and *CTT1* (right) using non-spatial (top) or spatial (bottom) analyses. All likelihoods values are relative to the most likely model under the corresponding analysis. To illustrate convergence, multiple MH chains have been run for each analysis using means only (red, > 1.2 × 10^6^ MH samples per chain), means and variances using the 𝒳^2^ or Wishart approach (blue, > 1.0 × 10^6^ MH samples per chain), means and variances using the extended moments approach (magenta, > 3.0 × 10^5^ MH samples per chain), or distributions (black, > 1.4 × 10^5^ MH samples per chain for non-spatial analyses and > 2.0 × 10^4^ MH samples per chain for spatial analyses).

**Figure S9:**
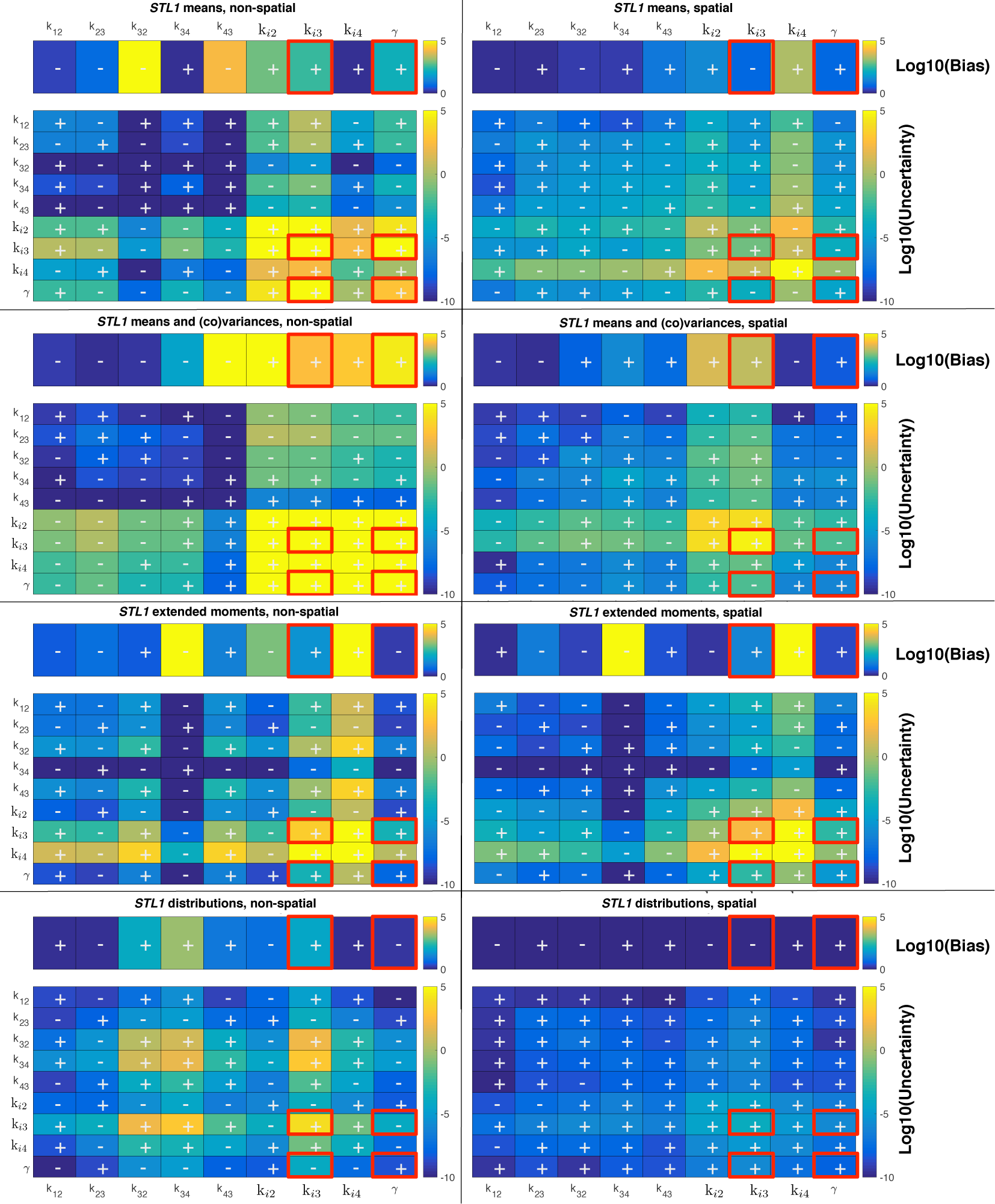
Bias and Uncertainty in the estimation of parameters for the *STL1* mRNA transcription. MCMC results are summarized for the means (top row), means and variances using the 𝒳^2^ or Wishart approach (second row), means and variances using the extended moments analysis (third row), or full FSP distributions (bottom row) using non-spatial (left) or spatial (right) models. For each of the eight panels, the top row illustrates the parameter estimation biases, where the colors denote the fold-change magnitude of the estimation error (in logarithmic scale). Blue boxes represent well estimated parameters and yellow boxes represent poorly estimated parameters. The signs in each box denote whether the parameter is over estimated (+) or underestimated (−). The matrices at the bottom of each panel illustrate the joint uncertainties in all parameter combinations. These are shown in a logarithmic scale from low variance (blue) to high variance (yellow). The signs in each box denote whether the corresponding parameter combination is positively or negatively correlated. The same color scale is used in all panels to illustrate the relative effects of uncertainty and bias for the different analyses. The entries boxed in red correspond to the parameter ellipses for *k*_*i*3_ and γ, which are shown in Fig. 4A.

**Figure S10:**
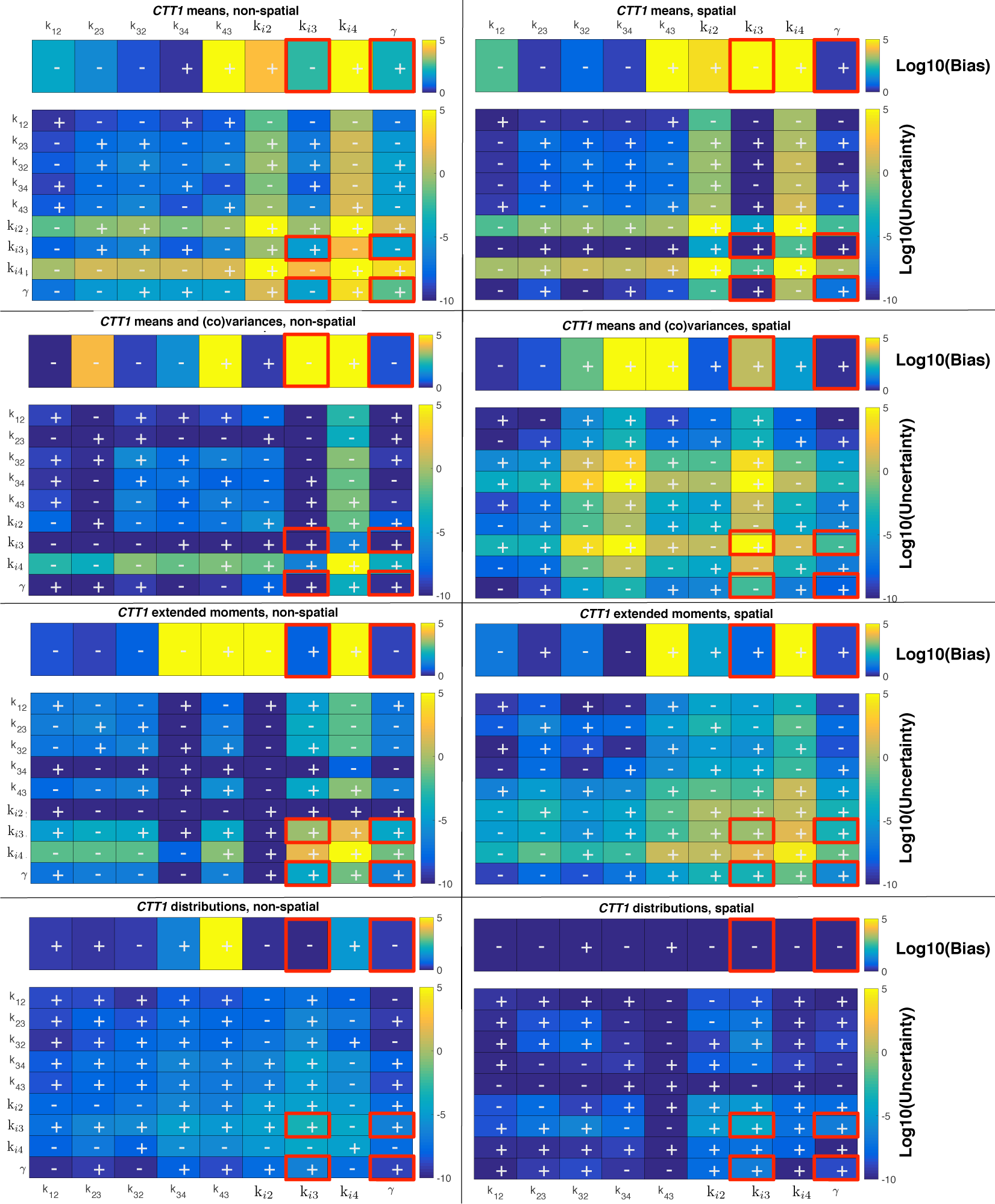
Bias and Uncertainty in the estimation of parameters for the CTT1 mRNA transcription. MCMC results are summarized for the means (top row), means and variances using the 𝒳^2^ or Wishart approach (second row), means and variances using the extended moments analysis (third row), or full FSP distributions (bottom row) using non-spatial (left) or spatial (right) models. For each panel, the top row illustrates the parameter estimation biases, where the colors denote the fold-change magnitude of the estimation error (in logarithmic scale). Blue boxes represent well estimated parameters and yellow boxes represent poorly estimated parameters. The signs in each box denote whether the parameter is over estimated (+) or underestimated (−). The matrices at the bottom of each panel illustrate the joint uncertainties in all parameter combinations. These are shown in logarithmic scale from low variance (blue) to high variance (yellow). The signs in each box denote whether the corresponding parameter combination is positively or negatively correlated. The same color scale is used in all six panels to illustrate the relative effects of uncertainty and bias for the six different analyses. The entries boxed in red correspond to the parameter ellipses for *k*_*i*3_ and γ, which are shown in Fig. S11A.

**Figure S11:**
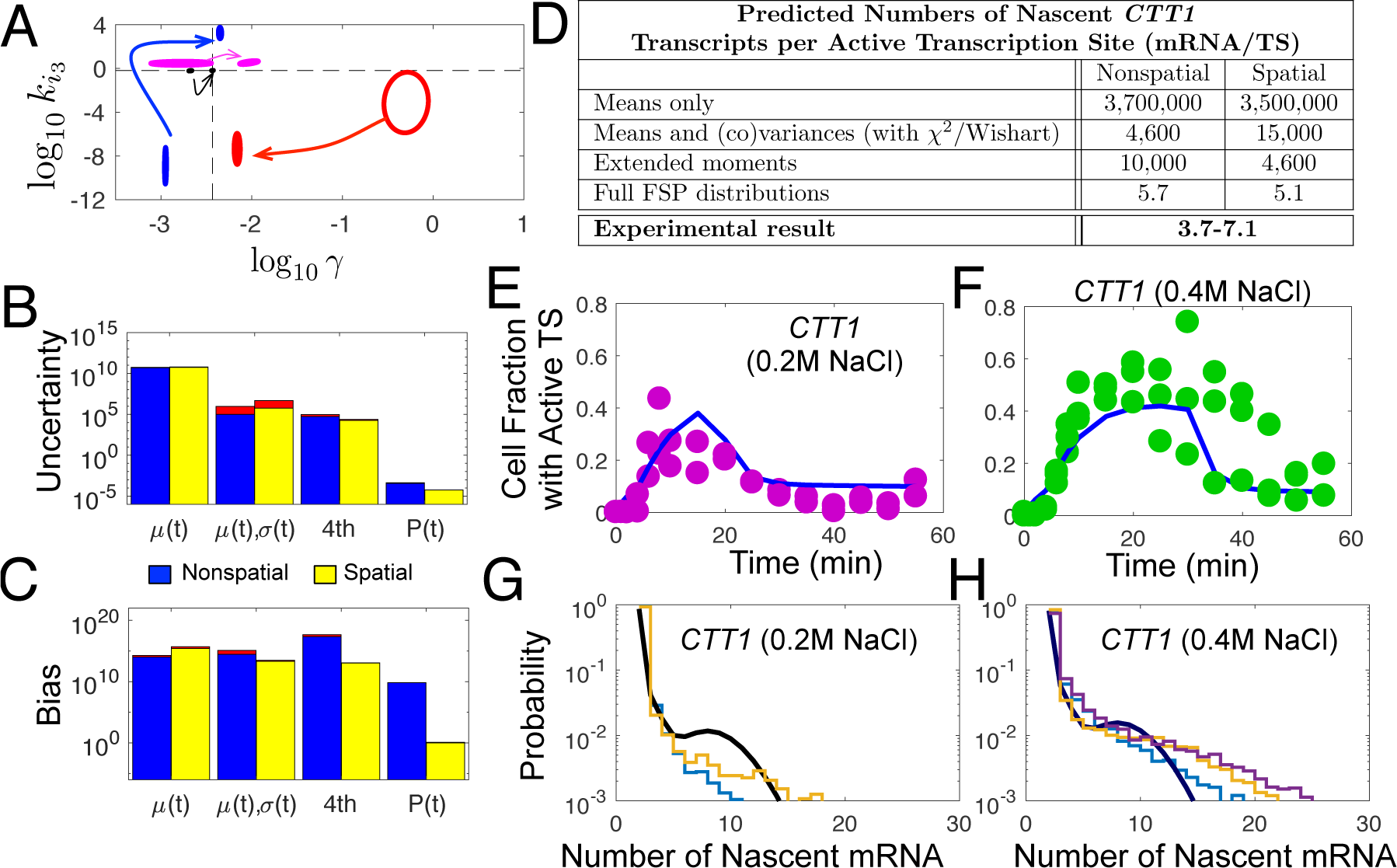
Overall bias and uncertainty quantification for CTT1 transcription regulation. (A) Ninety percent confidence ellipses for the degradation rate (γ) and the maximal transcription initiation rate (*k*_*i*3_) using the means only (*μ*(*t*), red), means and variances (*μ*(*t*), σ(*t*), blue), extended moment analyses (4th, magenta), or the full FSP distributions (**P**(*t*), black). Arrows show the effect of adding spatial information to the analysis. The dashed black lines show the baseline parameter combination corresponding to the best fit parameters for the spatial CTT1 model. (B) Total uncertainty in parameters for the four methods. (C) Total parameter bias for the four methods. For B and C, the results for non-spatial and spatial analyses are shown in blue and yellow, respectively, and the red regions show the convergence of two independent MHA samples. (D) Prediction for the average number of nascent mRNA per active CTT1 transcription site using each of the identified models. (E,F) Prediction for the fraction of cells with active CTT1 TS versus time at (E) 0.2M NaCl and (F) 0.4M NaCl osmotic shock. (G,H) Model fit (black) and biological replicas (blue, orange, purple) for the distributions of nascent *CTT1* mRNA per TS.

**Figure S12:**
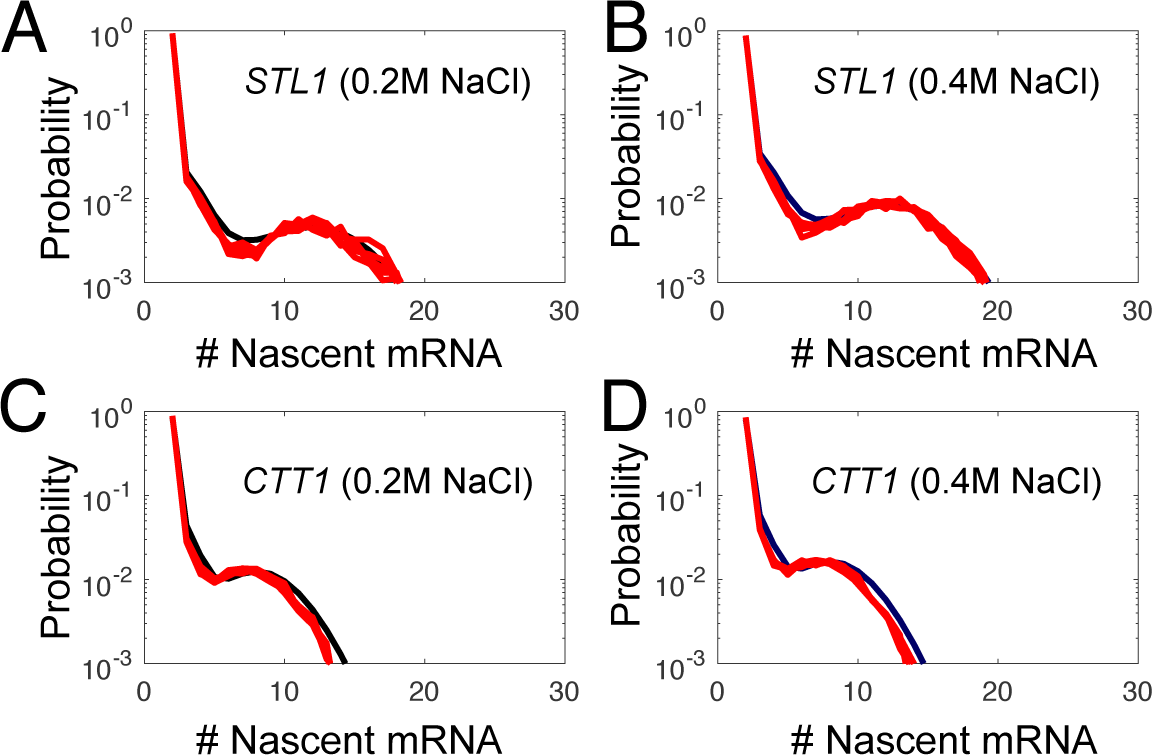
Comparison of FSP-based predictions (black) and several representative sets of 16,000 SSA simulations (red) for *STL1* TS intensity distributions following (A) 0.2M or (B) 0.4M NaCl osmotic shock and *CTT1* TS intensity distributions following (C) 0.2M or (D) 0.4M NaCl osmotic shock. The FSP and SSA analyses both use the best parameters found for the spatial FSP model (Table S3 for *STL1* and S4 for *CTT1*), with an elongation rate of 63nt/s.

**Table S1:** Locations for smRNA-FISH probes.

| ***CTT1* Probe Placement** |  | ***STL1* Probe Placement** |  |
| --- | --- | --- | --- |
| Probe # | 3' base relative to mRNA termination | Probe # | 3' base relative to mRNA termination |
| 1 | 1618 | 1 | 1630 |
| 2 | 1578 | 2 | 1604 |
| 3 | 1542 | 3 | 1577 |
| 4 | 1507 | 4 | 1546 |
| 5 | 1461 | 5 | 1518 |
| 6 | 1428 | 6 | 1493 |
| 7 | 1403 | 7 | 1449 |
| 8 | 1378 | 8 | 1389 |
| 9 | 1349 | 9 | 1364 |
| 10 | 1320 | 10 | 1325 |
| 11 | 1275 | 11 | 1282 |
| 12 | 1245 | 12 | 1250 |
| 13 | 1199 | 13 | 1200 |
| 14 | 1171 | 14 | 1172 |
| 15 | 1131 | 15 | 1144 |
| 16 | 1091 | 16 | 1104 |
| 17 | 1066 | 17 | 1078 |
| 18 | 1044 | 18 | 1035 |
| 19 | 1014 | 19 | 1001 |
| 20 | 987 | 20 | 971 |
| 21 | 953 | 21 | 913 |
| 22 | 913 | 22 | 884 |
| 23 | 884 | 23 | 856 |
| 24 | 854 | 24 | 830 |
| 25 | 832 | 25 | 805 |
| 26 | 810 | 26 | 768 |
| 27 | 788 | 27 | 743 |
| 28 | 743 | 28 | 702 |
| 29 | 693 | 29 | 670 |
| 30 | 671 | 30 | 635 |
| 31 | 648 | 31 | 610 |
| 32 | 615 | 32 | 574 |
| 33 | 591 | 33 | 545 |
| 34 | 568 | 34 | 509 |
| 35 | 546 | 35 | 472 |
| 36 | 519 | 36 | 435 |
| 37 | 491 | 37 | 396 |
| 38 | 468 | 38 | 356 |
| 39 | 445 | 39 | 324 |
| 40 | 390 | 40 | 287 |
| 41 | 313 | 41 | 250 |
| 42 | 289 | 42 | 223 |
| 43 | 237 | 43 | 184 |
| 44 | 166 | 44 | 138 |
| 45 | 105 | 45 | 113 |
| 46 | 57 | 46 | 86 |
| 47 | 27 | 47 | 59 |
| 48 | 0 | 48 | 24 |

**Table S2:** Parameterization of the Hog1p nuclear enrichment signal at 0.2M and 0.4M NaCl osmotic shock.

| Osmotic Shock Condition | Parameter Values |  |  |  |  |
| --- | --- | --- | --- | --- | --- |
| | $r_1$<br>($s^{-1}$) | $r_2$<br>($s^{-1}$) | $\eta$<br>- | $A$<br>- | $M$<br>- |
| 0.2M NaCl | $6.1 \times 10^{-3}$ | $6.9 \times 10^{-3}$ | 5.9 | $9.3 \times 10^9$ | $2.2 \times 10^{-2}$ |
| 0.4M NaCl | $6.1 \times 10^{-3}$ | $3.8 \times 10^{-3}$ | 5.9 | $9.3 \times 10^9$ | $2.2 \times 10^{-2}$ |

**Table S3:**
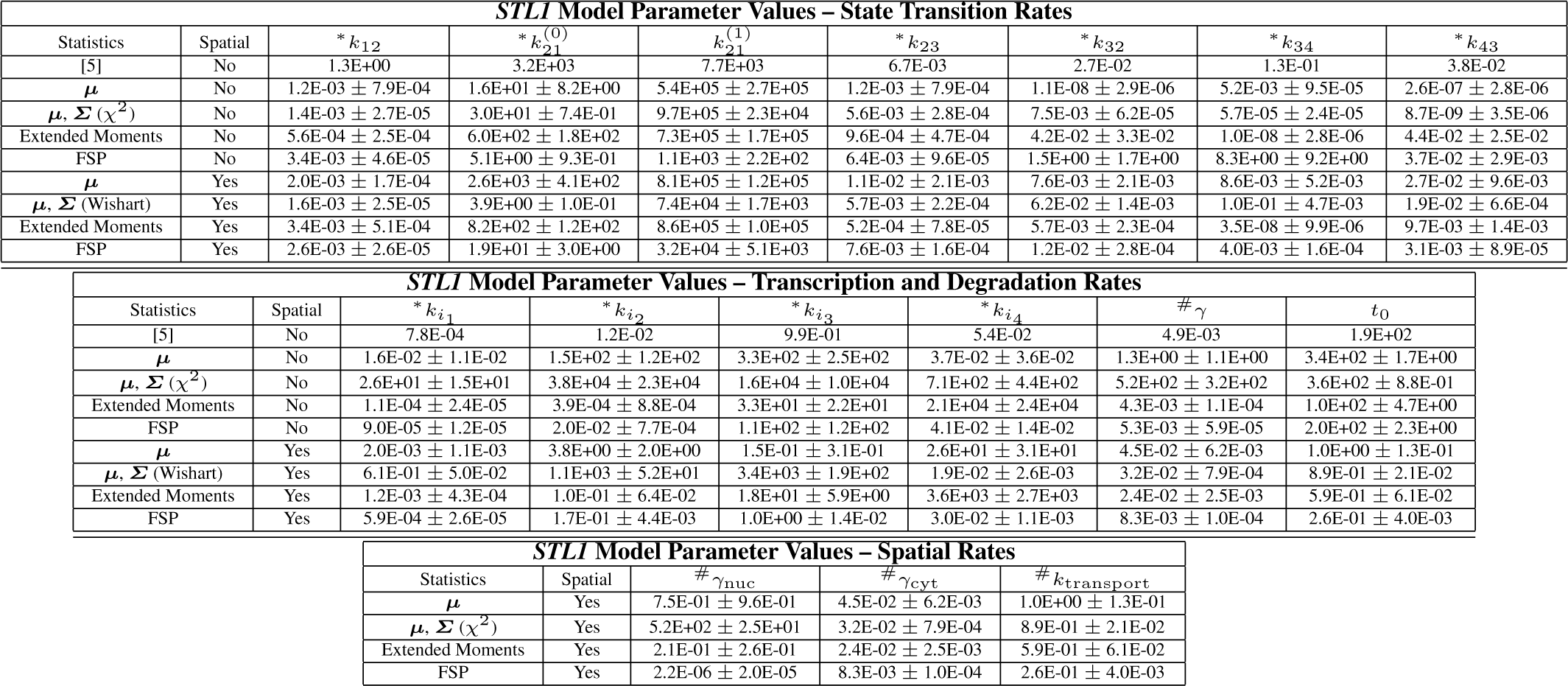
Parameter values (and uncertainty) for the final model for *STL1*. Each parameter set was identified using a different fluctuation analysis: means (μ), means and (co)variances (**μ** and ***σ/Σ*** with 𝒳^2^ or Wishart formulation), the extended moments-based analysis using the means and covariances and a likelihood function defined by the first four moments, or distributions; and a different assumption on the spatial fluctuations: non-spatial or spatial). Parameter uncertainties were identified using the Metropolis Hastings algorithm on the corresponding likelihood function and are listed as plus or minus one standard deviation. Units of all parameters denoted by (*) are s^−1^. Units of all parameters denoted by (^#^) are *Molecules*^−1^ *s*^−1^. Units of 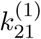 are *AUC*^−1^ *s*^−1^, where *AUC* is arbitrary units of Hog1p concentration. Units of *t*_0_ are *s*.

| **STLI Model Parameter Values – State Transition Rates** |  |  |  |  |  |  |  |  |
| --- | --- | --- | --- | --- | --- | --- | --- | --- |
| Statistics | Spatial | * $k_{12}$ | * $k_{21}^{(0)}$ | * $k_{21}^{(1)}$ | * $k_{23}$ | * $k_{32}$ | * $k_{34}$ | * $k_{43}$ |
| [5] | No | 1.3E+00 | 3.2E+03 | 7.7E+03 | 6.7E-03 | 2.7E-02 | 1.3E-01 | 3.8E-02 |
| $\mu$ | No | 1.2E-03 ± 7.9E-04 | 1.6E+01 ± 8.2E+00 | 5.4E+05 ± 2.7E+05 | 1.2E-03 ± 7.9E-04 | 1.1E-08 ± 2.9E-06 | 5.2E-03 ± 9.5E-05 | 2.6E-07 ± 2.8E-06 |
| $\mu, \Sigma (\chi^2)$ | No | 1.4E-03 ± 2.7E-05 | 3.0E+01 ± 7.4E-01 | 9.7E+05 ± 2.3E+04 | 5.6E-03 ± 2.8E-04 | 7.5E-03 ± 6.2E-05 | 5.7E-05 ± 2.4E-05 | 8.7E-09 ± 3.5E-06 |
| Extended Moments | No | 5.6E-04 ± 2.5E-04 | 6.0E+02 ± 1.8E+02 | 7.3E+05 ± 1.7E+05 | 9.6E-04 ± 4.7E-04 | 4.2E-02 ± 3.3E-02 | 1.0E-08 ± 2.8E-06 | 4.4E-02 ± 2.5E-02 |
| FSP | No | 3.4E-03 ± 4.6E-05 | 5.1E+00 ± 9.3E-01 | 1.1E+03 ± 2.2E+02 | 6.4E-03 ± 9.6E-05 | 1.5E+00 ± 1.7E+00 | 8.3E+00 ± 9.2E+00 | 3.7E-02 ± 2.9E-03 |
| $\mu$ | Yes | 2.0E-03 ± 1.7E-04 | 2.6E+03 ± 4.1E+02 | 8.1E+05 ± 1.2E+05 | 1.1E-02 ± 2.1E-03 | 7.6E-03 ± 2.1E-03 | 8.6E-03 ± 5.2E-03 | 2.7E-02 ± 9.6E-03 |
| $\mu, \Sigma$ (Wishart) | Yes | 1.6E-03 ± 2.5E-05 | 3.9E+00 ± 1.0E-01 | 7.4E+04 ± 1.7E+03 | 5.7E-03 ± 2.2E-04 | 6.2E-02 ± 1.4E-03 | 1.0E-01 ± 4.7E-03 | 1.9E-02 ± 6.6E-04 |
| Extended Moments | Yes | 3.4E-03 ± 5.1E-04 | 8.2E+02 ± 1.2E+02 | 8.6E+05 ± 1.0E+05 | 5.2E-04 ± 7.8E-05 | 5.7E-03 ± 2.3E-04 | 3.5E-08 ± 9.9E-06 | 9.7E-03 ± 1.4E-03 |
| FSP | Yes | 2.6E-03 ± 2.6E-05 | 1.9E+01 ± 3.0E+00 | 3.2E+04 ± 5.1E+03 | 7.6E-03 ± 1.6E-04 | 1.2E-02 ± 2.8E-04 | 4.0E-03 ± 1.6E-04 | 3.1E-03 ± 8.9E-05 |

| **STLI Model Parameter Values – Transcription and Degradation Rates** |  |  |  |  |  |  |  |
| --- | --- | --- | --- | --- | --- | --- | --- |
| Statistics | Spatial | * $k_{i1}$ | * $k_{i2}$ | * $k_{i3}$ | * $k_{i4}$ | # $\gamma$ | $t_0$ |
| [5] | No | 7.8E-04 | 1.2E-02 | 9.9E-01 | 5.4E-02 | 4.9E-03 | 1.9E+02 |
| $\mu$ | No | 1.6E-02 ± 1.1E-02 | 1.5E+02 ± 1.2E+02 | 3.3E+02 ± 2.5E+02 | 3.7E-02 ± 3.6E-02 | 1.3E+00 ± 1.1E+00 | 3.4E+02 ± 1.7E+00 |
| $\mu, \Sigma (\chi^2)$ | No | 2.6E+01 ± 1.5E+01 | 3.8E+04 ± 2.3E+04 | 1.6E+04 ± 1.0E+04 | 7.1E+02 ± 4.4E+02 | 5.2E+02 ± 3.2E+02 | 3.6E+02 ± 8.8E-01 |
| Extended Moments | No | 1.1E-04 ± 2.4E-05 | 3.9E-04 ± 8.8E-04 | 3.3E+01 ± 2.2E+01 | 2.1E+04 ± 2.4E+04 | 4.3E-03 ± 1.1E-04 | 1.0E+02 ± 4.7E+00 |
| FSP | No | 9.0E-05 ± 1.2E-05 | 2.0E-02 ± 7.7E-04 | 1.1E+02 ± 1.2E+02 | 4.1E-02 ± 1.4E-02 | 5.3E-03 ± 5.9E-05 | 2.0E+02 ± 2.3E+00 |
| $\mu$ | Yes | 2.0E-03 ± 1.1E-03 | 3.8E+00 ± 2.0E+00 | 1.5E-01 ± 3.1E-01 | 2.6E+01 ± 3.1E+01 | 4.5E-02 ± 6.2E-03 | 1.0E+00 ± 1.3E-01 |
| $\mu, \Sigma$ (Wishart) | Yes | 6.1E-01 ± 5.0E-02 | 1.1E+03 ± 5.2E+01 | 3.4E+03 ± 1.9E+02 | 1.9E-02 ± 2.6E-03 | 3.2E-02 ± 7.9E-04 | 8.9E-01 ± 2.1E-02 |
| Extended Moments | Yes | 1.2E-03 ± 4.3E-04 | 1.0E-01 ± 6.4E-02 | 1.8E+01 ± 5.9E+00 | 3.6E+03 ± 2.7E+03 | 2.4E-02 ± 2.5E-03 | 5.9E-01 ± 6.1E-02 |
| FSP | Yes | 5.9E-04 ± 2.6E-05 | 1.7E-01 ± 4.4E-03 | 1.0E+00 ± 1.4E-02 | 3.0E-02 ± 1.1E-03 | 8.3E-03 ± 1.0E-04 | 2.6E-01 ± 4.0E-03 |

Table S3: Parameter values (and uncertainty) for the final model for *STLI*.
| **STLI Model Parameter Values – Spatial Rates** |  |  |  |  |
| --- | --- | --- | --- | --- |
| Statistics | Spatial | # $\gamma_{nuc}$ | # $\gamma_{cyt}$ | # $k_{transport}$ |
| $\mu$ | Yes | 7.5E-01 ± 9.6E-01 | 4.5E-02 ± 6.2E-03 | 1.0E+00 ± 1.3E-01 |
| $\mu, \Sigma (\chi^2)$ | Yes | 5.2E+02 ± 2.5E+01 | 3.2E-02 ± 7.9E-04 | 8.9E-01 ± 2.1E-02 |
| Extended Moments | Yes | 2.1E-01 ± 2.6E-01 | 2.4E-02 ± 2.5E-03 | 5.9E-01 ± 6.1E-02 |
| FSP | Yes | 2.2E-06 ± 2.0E-05 | 8.3E-03 ± 1.0E-04 | 2.6E-01 ± 4.0E-03 |

**Table S4:**
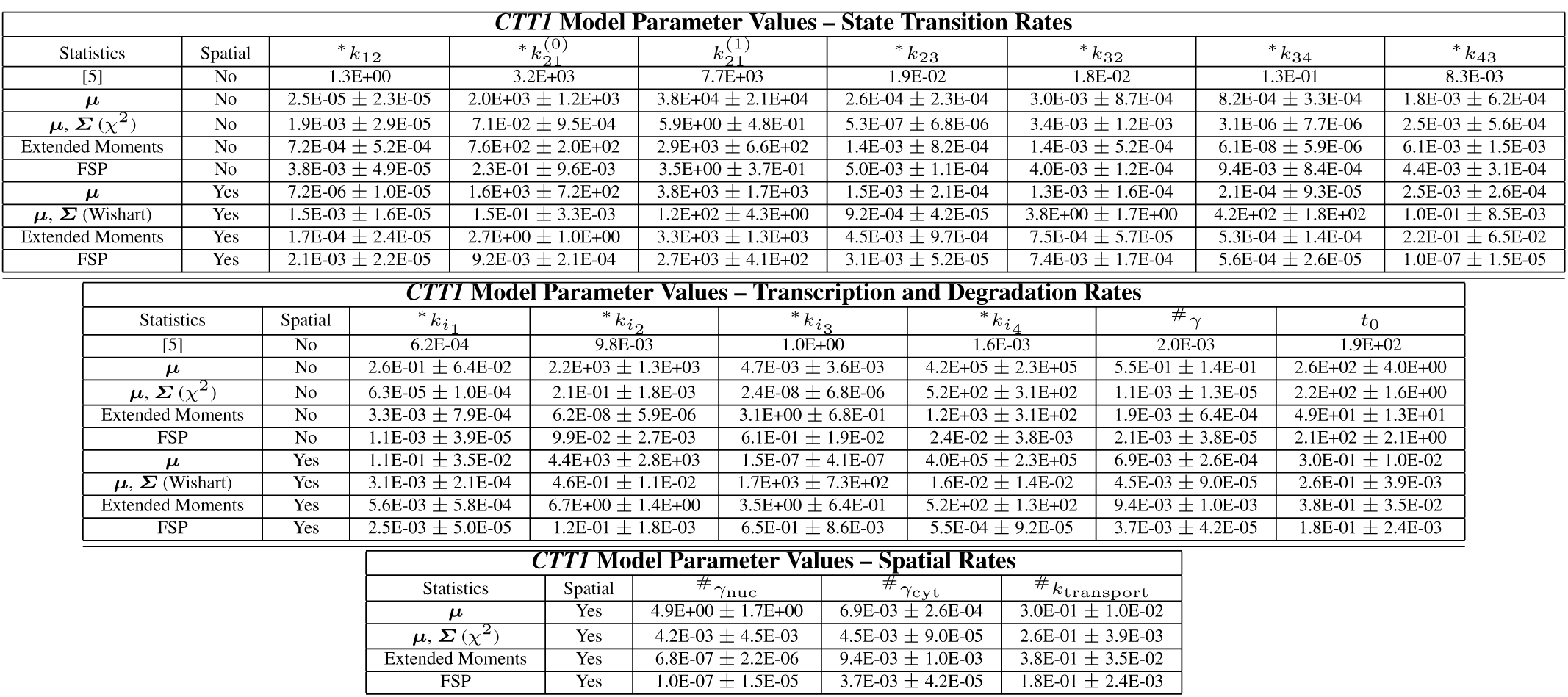
Parameter values (and uncertainty) for the final model for *CTT1.* Each parameter set was identified using a different fluctuation analysis: means (***μ***), means and (co)variances (***μ*** and ***σ/Σ*** with 𝒳^2^ or Wishart formulation), the extended moments-based analysis using the means and covariances and a likelihood function defined by the first four moments, or distributions computed with the FSP approach; and a different assumption on the spatial fluctuations: non-spatial or spatial). Parameter uncertainties were identified using the Metropolis Hastings algorithm on the corresponding likelihood function and are listed as plus or minus one standard deviation. Units of all parameters denoted by (*) are *s*^−1^. Units of all parameters denoted by (^#^) are *Molecules*^−1^ *s*^−1^. Units of 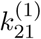 are *AUC*^−1^*s*^−1^, where *AUC* is arbitrary units of Hog1p concentration. Units of *t*_0_ are *s*.

**Table S5:** Relative log-likelihood values for different likelihoods that compare the means (***μ***), means and (co)variances (***μ*** and ***σ/Σ*** with 𝒳^2^ or Wishart formulation), the extended moments-based approach, or full distributions. Each row corresponds to a different combination of gene and identification strategy. Each column corresponds to a different likelihood function. Values presented are log_10_ of the actual likelihoods relative to the best value found for that objective function. For example, a value of zero states that the corresponding parameter set (Tables S4,S3) maximizes that likelihood function. A value –*x_i,j_*, in the *i*-row and *j*-column states that the parameters identified using the *i*^th^ likelihood yields 10^*x_ij_*^-fold worse fit to the *j*^th^ likelihood function. See Section 2.8 for detailed instructions on how to interpret this table. *The moment-based likelihood functions are approximated using the Central Limit Theorem as discussed in the Materials and Methods.

| **STLI Relative -Log<sub>10</sub>-Likelihood Values** |  |  |  |  |  |
| --- | --- | --- | --- | --- | --- |
|  |  | **Likelihood Function** |  |  |  |
| Fitting Data | Spatial | $\mu^*$ | $\mu$ and $\Sigma^*$ | Extended Moments* | Distributions |
| Means only | No | 0 | -1.72E+04 | -1.64E+04 | -6.97E+04 |
| Means and variances ($\chi^2$) | No | -4.46E+02 | 0 | -1.85E+04 | -6.26E+04 |
| Extended moments | No | -6.83E+02 | -3.71E+04 | 0 | -4.94E+04 |
| Full FSP Distributions | No | -2.75E+03 | -1.45E+04 | -6.64E+02 | 0 |
| Means only | Yes | 0 | -6.46E+04 | -7.32E+03 | -2.72E+04 |
| Means and variances (Wishart) | Yes | -7.62E+02 | 0 | -7.20E+03 | -1.77E+04 |
| Extended moments | Yes | -1.88E+03 | -4.75E+04 | 0 | -8.15E+04 |
| Full FSP Distributions | Yes | -1.03E+04 | -2.40E+04 | -3.45E+03 | 0 |
| **CTTI Relative -Log<sub>10</sub>-Likelihood Values** |  |  |  |  |  |
|  |  | **Likelihood Function** |  |  |  |
| Fitting Statistics | Spatial | $\mu^*$ | $\mu$ and $\Sigma^*$ | Extended Moments* | Distributions |
| Means only | No | 0 | -1.83E+05 | -1.48E+03 | -3.62E+05 |
| Means and variances ($\chi^2$) | No | -1.61E+03 | 0 | -2.29E+03 | -3.06E+04 |
| Extended moments | No | -9.49E+02 | -5.24E+04 | 0 | -9.39E+04 |
| Full FSP Distributions | No | -4.76E+03 | -5.62E+03 | -1.83E+03 | 0 |
| Means only | Yes | 0 | -3.85E+05 | -1.94E+03 | -3.57E+05 |
| Means and variances (Wishart) | Yes | -3.11E+03 | 0 | -1.94E+04 | -2.45E+04 |
| Extended moments | Yes | -4.13E+03 | -1.25E+05 | 0 | -2.36E+05 |
| Full FSP Distributions | Yes | -1.56E+04 | -1.60E+04 | -6.65E+03 | 0 |

**Table S6:** Elongation rates identified for *CTT1* transcription using the best parameters identified in Tables S4 and the measured TS intensity data. For each case – means (***μ***), means and (co)variances (***μ and Σ***), extended moments, or distributions – we present the upper bound on the rates using the simplified theoretical model and the more precise rates using the detailed FSP-TS analysis, when possible. Uncertainty in these rates is given as the standard error of the mean computed from the five biological replicates. (*Figs. 1D, 4D, and S11 use the rate identified from *CTT1* TS intensity measurements and the corresponding fit of the elongation rate with the spatial FSP analysis.)

| **Estimation of *CTT1* Transcription Elongation Rates (Nt/s)** |  |  |  |
| --- | --- | --- | --- |
| **Analysis** | **Spatial** | **Simplified Theory** | **Detailed FSP** |
| Means only | No | 5.93E+07 ± 5.9E+06 | N/A |
| Means and variances ($\chi^2$) | No | 7.25E+04 ± 7.3E+03 | N/A |
| Extended moments | No | 1.65E+05 ± 1.7E+04 | N/A |
| Full FSP Distributions | No | 85.8 ± 8.6 | 45 ± 14 |
| Means only | Yes | 5.54E+07 ± 5.6E+06 | N/A |
| Means and (co)variances (Wishart) | Yes | 2.39E+05 ± 2.4E+04 | N/A |
| Extended moments | Yes | 7.27E+04 ± 7.3E+03 | N/A |
| Full FSP Distributions | Yes | 91.2 ± 9.1 | **\*63 ± 13** |

### Footnotes

1 A GitHub download page will be created upon acceptance of this manuscript.

2 The time offset parameter *t*_0_ is only necessary for the non-spatial model and is identified as nearly zero for the spatial model.

